# RamiGlyph Captures Microglial Morphological Diversity and Predicts Functional States in Ischemia Reperfusion and Amyloid Pathology

**DOI:** 10.64898/2026.09.10.750561

**Authors:** Yuan Qu, Yueming Xiao, Tian Lan, Jizhuang Xu, Jinliang Liu, Qiuting Qian, Junru Liu, Ying Chi

**Affiliations:** Department of Pharmacy of the Second Affiliated Hospital of Zhejiang University School of Medicine, and Zhejiang University-University of Edinburgh Institute (ZJE), Zhejiang University, No.866 Yu Hang Tang road, 310058, Zhejiang Province, China; Edinburgh Medical School, College of Medicine and Veterinary Medicine, The University of Edinburgh, Edinburgh, United Kingdom; School of Pharmaceutical Sciences, Wenzhou Medical University, Chashan Higher Education 17 Zone, 325035, Zhejiang Province, China; College of Medicine, Jiaxing University, Jiaxing, Zhejiang Province, China

## Abstract

Microglia are resident immune cells of the central nervous system, whose ramified processes rapidly remodel in response to injury. However, how to capture subtle morphological changes and whether functional state can be predicted from morphology remain open questions. Here, we present RamiGlyph, a contrastive learning framework integrating topological and structural features, trained on more than 20,000 reconstructed microglia. RamiGlyph not only distinguishes physiological and pathological states of microglia but also generalizes to neuronal cell type classification. Projection of microglia morphological embeddings revealed a continuum rather than discrete classes, from which a morphology score was derived to quantify dynamic process remodeling. To link morphology with function, Gromov Wasserstein optimal transport was used to align unpaired morphological and functional data across stages of ischemia reperfusion injury and amyloid pathology. These alignment results enable prediction of microglial functional states using RamiGlyph embeddings alone, with prediction reliability increasing upon cell aggregation. In summary, RamiGlyph provides a robust framework for resolving continuous microglial morphological variation and linking morphology to functional states across acute and chronic neuropathological contexts.

## Introduction

Microglia continuously remodel their processes during homeostasis and in response to perturbation^1^. Within minutes, process extension and retraction support environmental surveillance and rapid adaptation to injury, including synaptic remodeling and inflammatory responses, among other functions^2,3^. These rapid structural changes make morphology a sensitive phenotypic readout of microglial state that can be measured at single cell resolution while preserving anatomical context. However, how to accurately capture subtle morphological variation and whether functional state can be reliably predicted from morphology remain open questions.

Traditionally, ramified microglia have been associated with homeostatic surveillance, whereas amoeboid microglia have been linked to immune activation^4,5^. However, single cell transcriptomics has revealed a diverse continuum of microglial states^6^, challenging such simple associations between morphology and function states ^7^. This gap is further amplified by conventional morphological quantification, which often relies on two-dimensional images and a few predefined morphological features such as Sholl profiles, branch counts and process length^8,9^. Such features are sensitive to imaging and sampling differences and ignore three-dimensional (3D) organization, including process orientation and branching geometry. Consequently, current morphological phenotyping remains incomplete, difficult to standardize and insufficient to resolve the full diversity of microglial states.

Data driven approaches have begun to address these limitations. MorphOMICs derived a persistence image from each microglial reconstruction to encode process topology and compared bootstrapped population averages across conditions without user selected morphometric features^10^. The method, however, relied on predefined features rather than learning the representation directly from data. Learned representations have advanced faster for neurons^11^. MorphVAE sampled soma to terminal walks from reconstructed arbors and encoded them using a variational sequence-to-sequence autoencoder^12^. TreeMoCo used a Tree-LSTM encoder with morphology specific augmentations and contrastive learning to obtain self-supervised embeddings^13^. MorphoGNN represented neuronal arbors as graphs and learns point-level structural features through neighborhood aggregation^14^. SGTMorph combined graph neural networks and Transformers to capture local topology and global structural relationships^15^. Most existing morphology representation methods were trained on neuronal datasets and may not capture the dynamic remodeling of microglial processes, highlighting the need for a microglia-specific framework.

A detailed description of morphology alone may not be sufficient to infer functional state. Many studies integrate morphological analysis with immunostaining-based molecular profiling, but these approaches are typically limited to a small set of predefined markers^16,17^. Transcriptomics provides a broader view of microglial molecular states, yet integrating it with morphology remains difficult. Morphological reconstruction requires intact cellular processes, whereas single cell RNA sequencing usually requires tissue dissociation. Patch-seq enables paired morphological and transcriptomic measurements^18^, but microglia are highly sensitive to mechanical and purinergic stimuli, which can rapidly remodel their processes^19^. Thus, reliably linking morphological and transcriptomic states remains a major challenge.

Here we develop a self-supervised graph learning framework called RamiGlyph that encodes 3D microglial reconstructions through complementary topological and structure branches. Trained on 23,637 unlabeled cells, the model learns transferable embeddings that outperform existing methods. Microglial morphology embeddings formed a continuum, from which we derived a process complexity score for cross-dataset validation. We then use Gromov-Wasserstein optimal transport (GW-OT) to align morphology with single cell transcriptomic activity in Ischemia reperfusion and amyloid pathology ^20,21^. Overall, morphology carries predictive information for eight microglial functional states, with limited single cell accuracy but improved population level reliability.

## Results

### RamiGlyph architecture: self-supervised dual-branch framework for microglial morphology representation

Conventional microglial morphometry relies largely on hand crafted descriptors, which can limit scalability and introduce operator dependent bias. To address these constraints, we developed RamiGlyph, a self-supervised framework that learns morphological representations directly from 3D morphology without predefined morphometric features.

RamiGlyph comprises four modules: graph construction, dual-view augmentation, dual branch encoding and self-supervised prototype learning. Each 3D reconstruction was converted from an SWC file into a soma rooted tree graph. Sampling points served as nodes and parent child relationships as edges, preserving the branching topology. Node features included coordinates, radius, node type, graph distance from the soma, branch level, branch identity and random walk positional encoding (Fig. 1A).

**Fig. 1.**
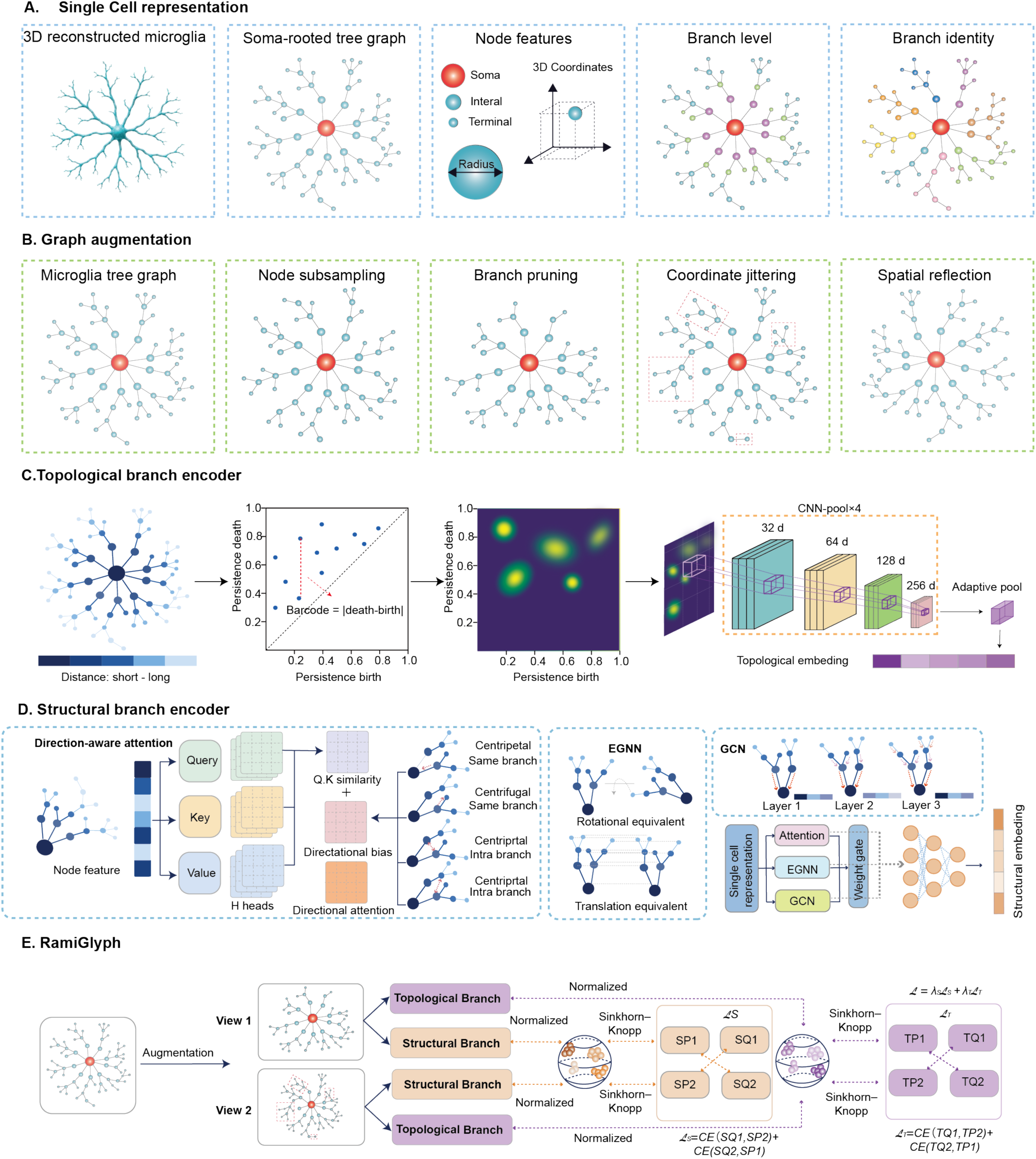
RamiGlyph architecture. **A** Graph construction. Each SWC reconstruction is converted into a soma-rooted attributed graph, with sampling points as nodes and parent–child relationships as bidirectional edges. Each node carries its 3D coordinates, radius and node type, together with three soma-referenced descriptors: soma distance, branch level and primary branch identity. **B** Dual-view augmentation. Two views are generated from each graph by node subsampling, branch pruning, coordinate jittering and spatial reflection. The soma and all branch points are preserved, so that branching topology remains intact. **C** Topological branch. Zero-dimensional extended persistent homology is computed with shortest-path distance from the soma as the filtration value, yielding a persistence diagram that is converted into a persistence image and encoded by a convolutional network. **D** Structural branch. Each block processes the same node representation through three parallel pathways. Direction-aware attention distinguishes centripetal, centrifugal, intra branch and inter branch node pairs, an E(n)-equivariant network incorporates 3D distances, and local graph convolution operates along SWC-defined edges. The three outputs are combined by a node-wise gating network. **E** Self-supervised prototype learning. Normalized embeddings from the two views are assigned to a set of learnable prototypes, balanced by the Sinkhorn–Knopp algorithm; the assignment of one view supervises the prototype prediction of the other.

For self-supervised training, RamiGlyph generates two augmented views of each cell graph through node subsampling, branch pruning, coordinate jittering, translation and spatial reflection (Fig. 1B). The soma and graph connectivity are preserved, so that augmentation perturbs local detail without altering cell identity. The resulting representations therefore reflect branching structure rather than reconstruction differences.

Each augmented view is encoded by a dual branch network that captures complementary structural and topological features. The topological branch summarizes the global organization of the process by zero-dimensional extended persistent homology, filtered by graph distance from the soma (Fig. 1C). The resulting persistence diagram is converted into a persistence image and encoded by a lightweight convolutional network to yield a cell-level topological embedding.

The structural branch encodes node and edge features through stacked parallel blocks. Each block combines gated graph convolution^22^, equivariant graph neural networks and direction aware attention to capture local connectivity^23^, 3D geometry and soma referenced directionality-centripetal, centrifugal and inter-branch. At each node, the pathway outputs are fused by a learned gating mechanism and refined by a multilayer perceptron. Global mean pooling then aggregates these node representations into a cell level structural embedding (Fig. 1D).

Training organizes the representation space using learnable morphological prototypes, without manual annotation. The structural and topological branches maintain separate prototype sets. Within each branch, normalized embeddings from the two augmented views are softly assigned to prototypes, with the Sinkhorn-Knopp algorithm enforcing balanced assignments to prevent collapse^24^. The assignment of one view then serves as the prediction target for the other, and vice versa. This swapped prediction objective yields augmentation invariant representations with an intrinsic clustering structure, learned without labels or negative samples^24^. The final loss is a weighted combination of the structural and topological terms (Fig. 1E).

The two branches yield complementary embeddings capturing local geometry and global topology, which are concatenated and projected into a unified single cell representation for downstream analysis.

### RamiGlyph outperforms existing morphology models through topological and direction-aware encoding

We then compared RamiGlyph with representative morphology-representation methods developed mainly for neuronal and tree like structures, including SGTMorph, MorphoGNN, Tree-MoCo and MorphVAE^12–15^. MorphOMICs and morphometric features, both widely used for microglial morphology, were included as additional baselines ^8,10^. We further combined RamiGlyph with morphometric features to test whether learned and classical representations capture complementary aspects of cell morphology.

Each model was trained on the same training set, with the loss decreasing and stabilizing in all cases (Supplementary Fig. 1A). We then evaluated the resulting embeddings by k nearest neighbour (KNN) classification on an independent validation set, inferring each cell’s experimental condition and brain-region label from its nearest training set neighbours in the embedding space.

With K = 5, RamiGlyph reached an average macro-F1 of 0.52 and accuracy of 0.59 across six datasets, exceeding morphometric features, MorphOMICs and other deep-learning baselines^10^. Combining RamiGlyph with morphometric features raised these to 0.65 and 0.70, indicating that the two representations carry complementary information (Fig. 2A, Supplementary Fig. 1B). The ranking was stable across values of K (Fig. 2A, Supplementary Fig. 1B).

**Fig. 2.**
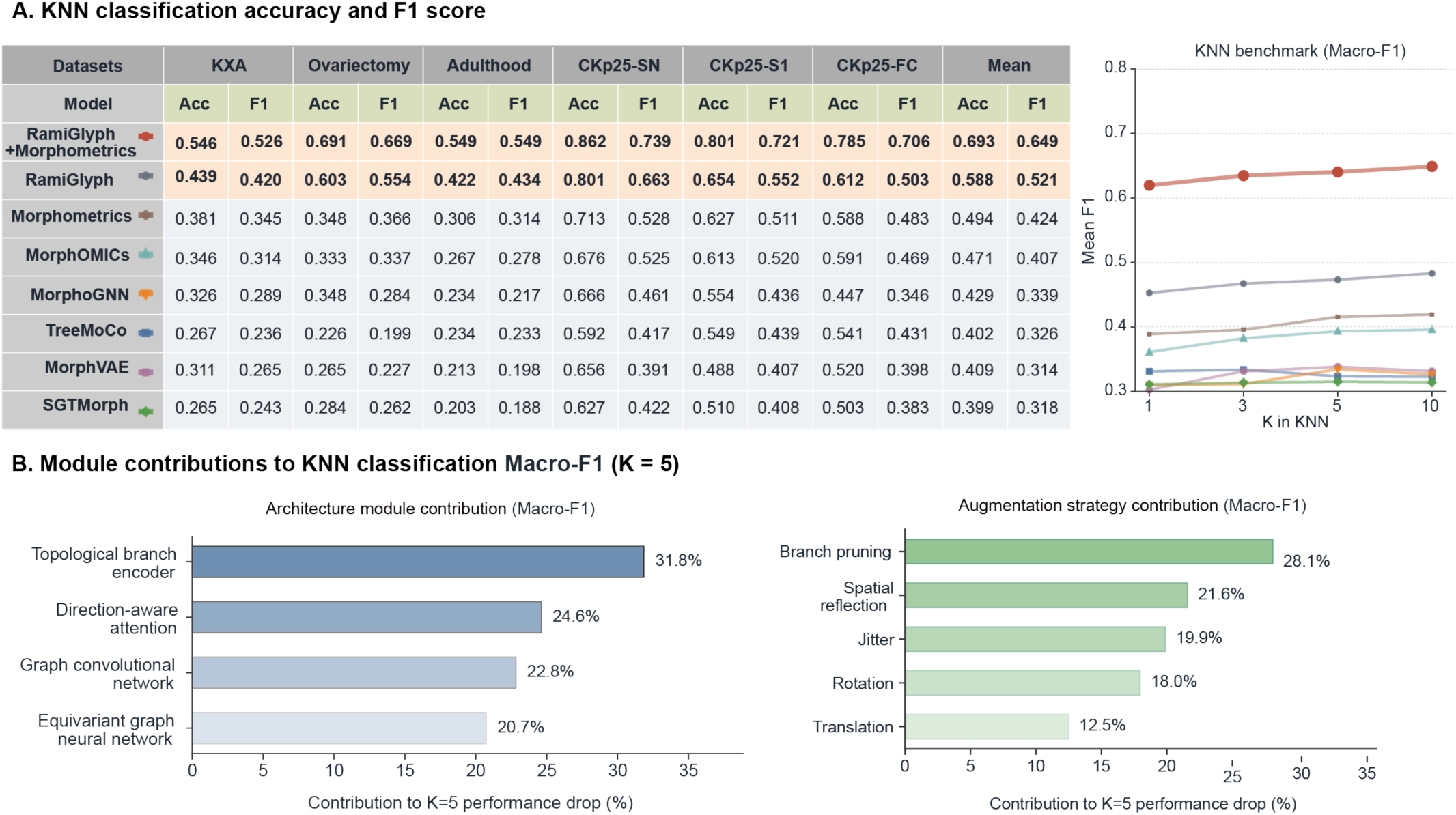
RamiGlyph improves KNN classification and reveals module contributions. **A** KNN performance at K = 5 for RamiGlyph, RamiGlyph with Morphometrics, and six representative morphology models across six datasets, reported as accuracy (ACC) and macro F1. The line plot shows mean macro F1 across ten datasets for K = 1, 3, 5 and 10. **B** Ablation analysis of module contributions to KNN performance at K = 5. Bars show the mean macro F1 decrease after removing each module or augmentation strategy across ten datasets.

Ablation of individual modules identified the topological branch encoder as the largest contributor, reducing average macro-F1 by 0.12 and accounting for 33.0% of the total effect (Fig. 2B). Direction aware attention, graph convolution and the equivariant graph neural network contributed 24.70%, 21.90% and 20.50% (Fig. 2B). Accuracy followed the same ranking, indicating that topological hierarchy and direction aware modelling are central to the learned representations (Supplementary Fig. 1C).

Ablation of the augmentations identified branch pruning as the most influential, reducing average macro-F1 by 0.10 and accounting for 28.10% of the total effect. Spatial reflection, jitter, rotation and translation contributed 21.60%, 19.90%, 18.00% and 12.50% (Fig. 2B). Accuracy followed the same ranking, indicating that that morphology preserving perturbations, particularly local branch pruning, provide the strongest augmentation signal (Supplementary Fig. 1C).

RamiGlyph therefore outperforms existing morphology representation methods using a KNN evaluation protocol established in previous studies^13–15^. Ablation attributes this advantage mainly to topological branch encoding, direction aware attention and branch level augmentation.

### RamiGlyph embeddings distinguish biological conditions at population and single-cell levels

This raised the question of whether the learned embeddings capture biologically meaningful differences in microglial morphology. Because individual microglia are morphologically heterogeneous, we first used bootstrap analysis to test whether the embeddings separate biological conditions at the population level, including brain regions, hormonal and pharmacological perturbations, and disease stages^25^. We then examined whether this structure persisted at single-cell resolution (Fig. 3A).

**Fig. 3.**
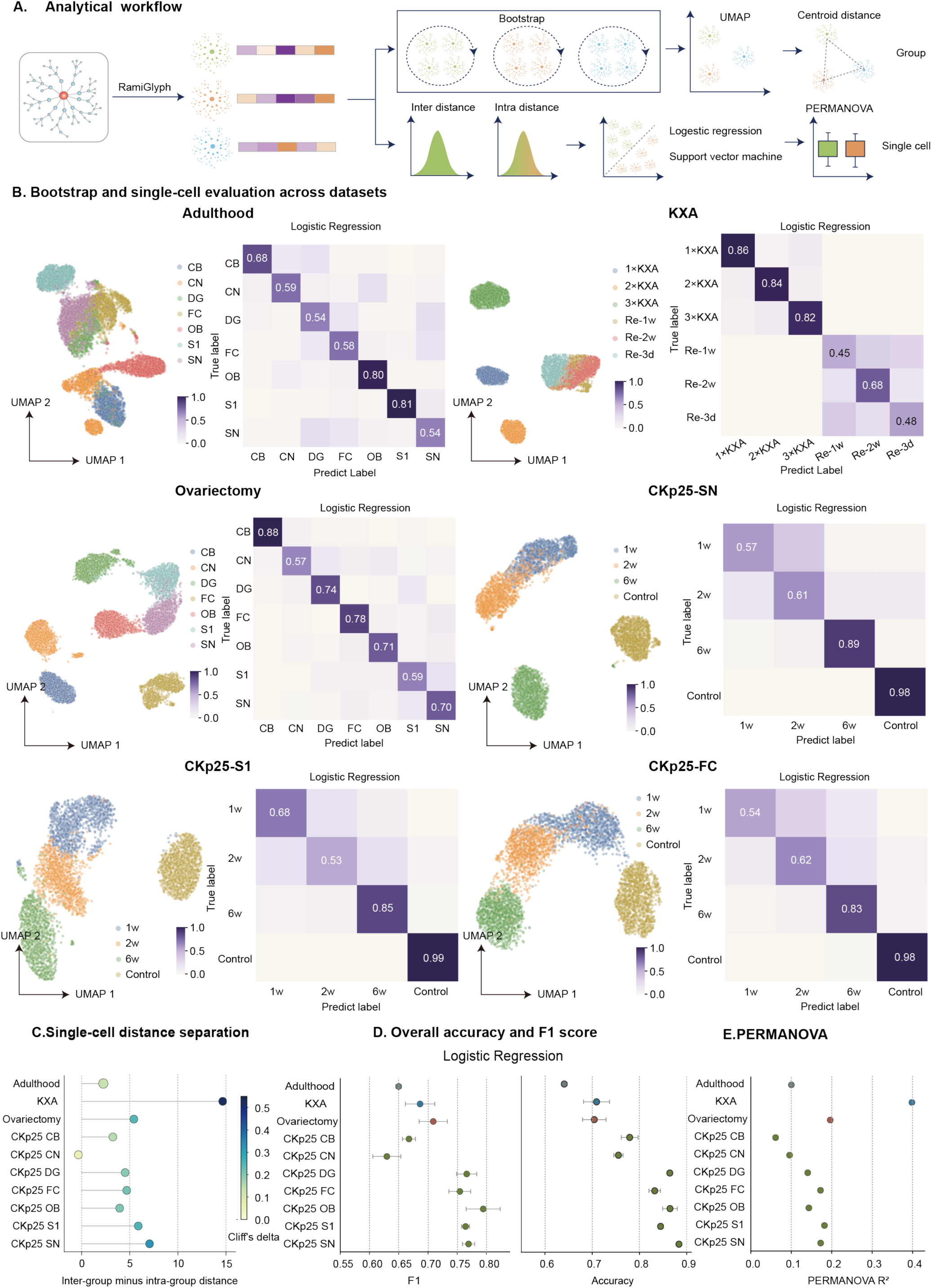
RamiGlyph embeddings capture population- and single-cell-level microglial morphological heterogeneity. **A.** Analysis workflow. Embeddings were aggregated into bootstrapped pseudo-populations for UMAP visualization and centroid distance calculation. At the single-cell level, within-group and between-group distances were compared, linear classifiers (logistic regression and linear SVM) were trained, and PERMANOVA was used to test multivariate separation. **B**. UMAP projections of bootstrap aggregated embeddings and logistic regression confusion matrices for the Adulthood, KXA and Ovariectomy datasets and representative CKp25 regions (SN, S1 and FC). For each group, 1,000 bootstrap embeddings were generated by resampling 15 cells with replacement. Numbers on the diagonal indicate classification accuracy. SN, substantia nigra; S1, primary somatosensory cortex; FC, frontal cortex; Re, recovery; w, week. **C.** Single cell distance separation across datasets, quantified as the difference between the mean inter group and intra group Euclidean distances in the embedding space. Point color indicates the corresponding Cliff’s delta effect size; positive values indicate greater separation between biological groups than within groups. CB, cerebellum; CN, caudate nucleus; DG, dentate gyrus; OB, olfactory bulb. **D.** Macro-F1 score and overall accuracy of logistic regression classifiers across datasets. Points and error bars represent the mean ± s.d. over five stratified cross-validation folds. **E.** Proportion of embedding space variation explained by biological group labels, quantified by PERMANOVA R² with 999 permutations. Higher R² values indicate stronger separation among biological groups.

Bootstrap analysis showed that distinct biological states formed reproducible population level structures in the embedding space. UMAP visualization and centroid distances in the 512-dimensional space gave consistent results across all four datasets. In Adulthood, embeddings separated by brain region, with the largest distance between cerebellum (CB) and primary somatosensory cortex (S1) and the smallest between dentate gyrus (DG) and substantia nigra (SN) (Fig. 3B, Supplementary Fig.3). In ketamine xylazine acepromazine (KXA) anesthesia, acute drug perturbation was clearly distinguished from recovery with centroid distances of 54.77 between 3×KXA and Recovery-2w but only 4.00 between Recovery-1w and Recovery-2w (Fig. 3B, Supplementary Fig.3). In Ovariectomy, region-associated separation persisted under altered hormonal status, with CB and frontal cortex (FC) most distinct (Fig. 3B, Supplementary Fig.3). In CKp25, disease time points separated from controls in a region dependent manner, most clearly in SN, S1 and FC (Fig. 3B, Supplementary Fig.2, 3).

Consistent with these distances, permutational multivariate analysis of variance (PERMANOVA) showed that group labels explained a measurable fraction of variance in the embedding space, with R^2^ highest in KXA (0.40), intermediate in Ovariectomy (0.20) and lowest in Adulthood (0.10)^26^. Across CKp25 brain regions, R^2^ ranged from 0.06 to 0.18, with the highest values in S1, SN and FC (Fig. 3E).

To test whether this organization was reflected at the level of individual cell-to-cell distances, we compared embedding distances within and between groups. Inter-group distances exceeded intra-group distances in most datasets (Fig. 3C). Separation was strongest in KXA, where the mean inter-group distance exceeded the intra-group distance by 14.66 (Cliff’s δ = 0.52), followed by Ovariectomy (5.43, δ = 0.25) and Adulthood (2.74, δ = 0.12), consistent with brain-region identity contributing less than perturbation or disease status. Within CKp25, separation was clearest in SN (7.06, δ = 0.33) and S1, whereas cochlear nucleus (CN) approached zero, indicating weaker morphological separation or greater within-region heterogeneity (Fig. 3C).

We then trained linear classifiers to predict each cell’s group from its embedding alone. Logistic regression achieved macro-F1 scores of 0.65, 0.69 and 0.71 in Adulthood, KXA and Ovariectomy (Fig. 3B, D). Performance was higher across CKp25 brain regions (0.75–0.80 in DG, FC, OB, S1 and SN), suggesting that disease time point was more readily predictable in these regions. A linear support vector machine (SVM) gave broadly consistent, slightly lower scores, indicating that the structure was not classifier specific (Supplementary Fig.4)^27^.

**Fig. 4.**
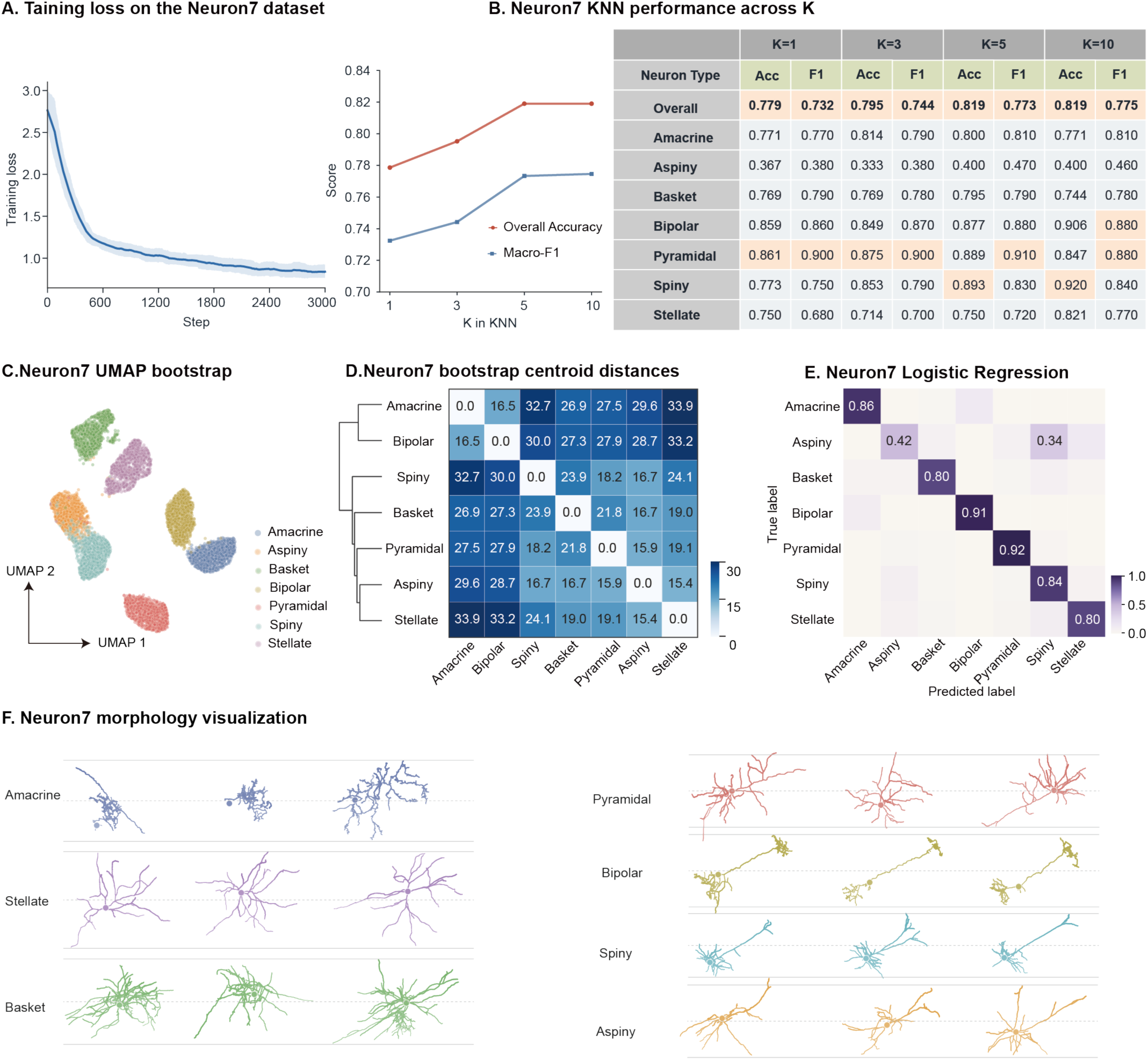
RamiGlyph generalizes to neuronal morphology classification in the Neuron7 dataset. **A** Training loss during self-supervised learning on Neuron7. **B** KNN classification accuracy (ACC) and macro F1 for K = 1, 3, 5 and 10. Validation neurons were classified using their nearest neighbours in the training embedding space. **C** UMAP projection of bootstrap aggregated embeddings for seven neuronal classes. For each class, 1,000 bootstrap embeddings were generated by resampling 15 neurons with replacement. **D** Heatmap of pairwise Euclidean distances between class centroids. The dendrogram shows average linkage hierarchical clustering, with larger distances indicating greater morphological separation. **E** Confusion matrix from five-fold stratified cross validated multinomial logistic regression on single neuron embeddings. Numbers on the diagonal indicate classification accuracy. **F** Representative morphology visualizations of the three neurons nearest to the centroid of each class in the embedding space.

The embeddings separated brain regions, perturbations and disease states consistently at the population, pairwise and single-cell levels, providing a quantitative representation of microglial morphology.

### RamiGlyph captures neuronal morphology and transfers without retraining

To test whether RamiGlyph generalizes to other tree-like cell types, we applied it to Neuron7, a dataset comprising seven neuronal types (Supplementary Fig.5A). Training converged stably (Fig. 4A). A KNN classifier on held-out cells achieved an accuracy of 0.82 and a macro-F1 score of 0.77 with k=5, while logistic regression and a linear SVM performed similarly, achieving accuracies of 0.84 and 0.83 and macro-F1 scores of 0.79 and 0.78, respectively (Fig. 4B, E; Supplementary Fig. 5B). Bootstrap UMAP and centroid distance analyses showed that the seven types formed distinguishable class related distributions in the embedding space (Fig. 4C, D). Visualizing the three cells closest to each type centroid revealed consistent morphologies within types, providing visual confirmation that embedding proximity reflects morphological similarity (Fig. 4F).

**Fig. 5.**
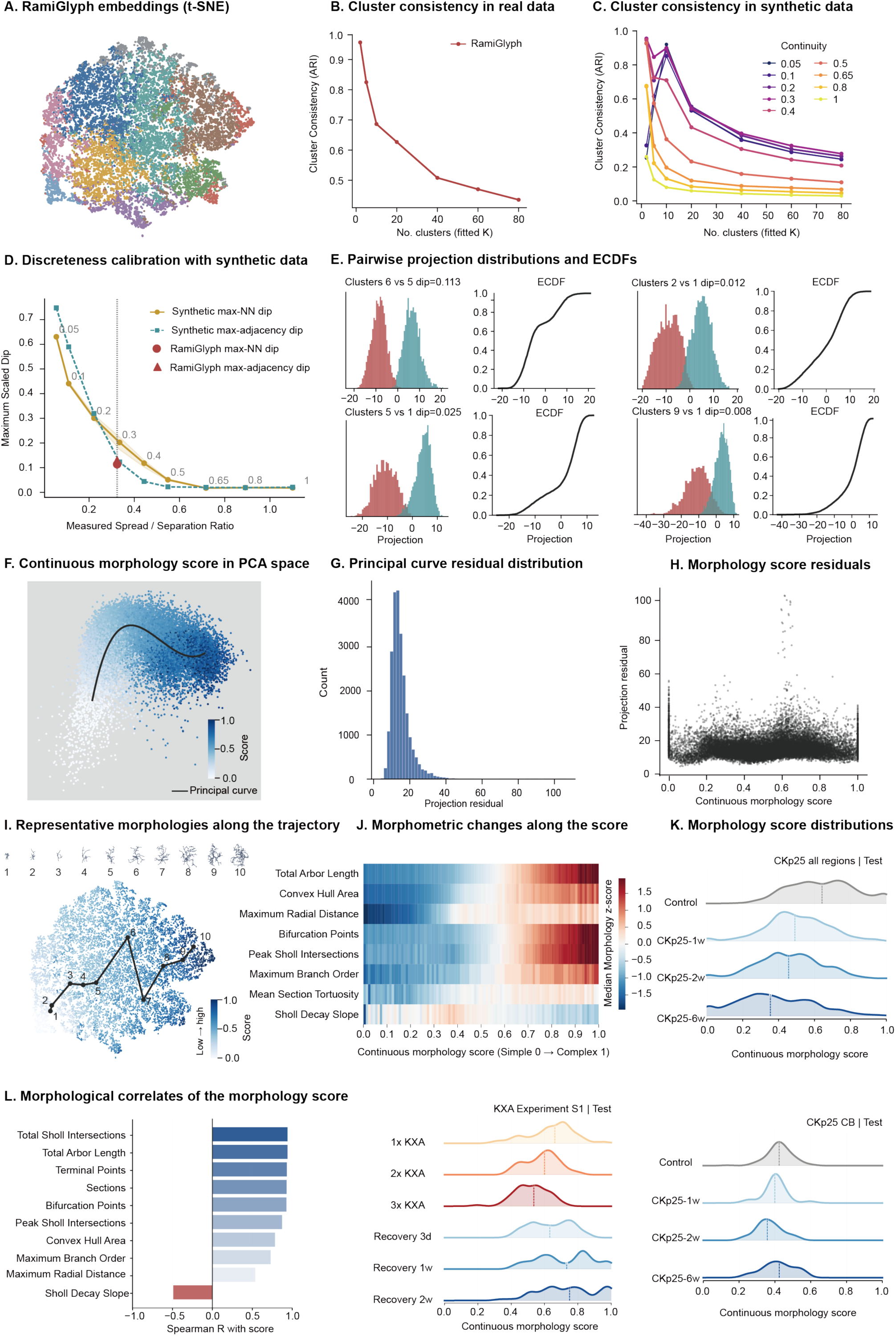
Continuous organization and morphometric interpretation of the RamiGlyp hembedding. **A** t-SNE visualization of RamiGlyph embeddings colored by Gaussian mixture model clusters. **B** Cluster consistency across fitted cluster numbers (K), quantified by the mean pairwise Adjusted Rand Index (ARI) across repeated fits. **C** ARI profiles of synthetic reference datasets with different degrees of continuity. **D** Calibration of morphological continuity using synthetic datasets with different levels of separation. The spread to separation ratio reflects the overlap between neighboring groups, while NN dip and adjacency dip describe separation based on local neighborhoods and overall connectivity, respectively. The maximum scaled dip represents the strongest local evidence of discreteness. Comparison with synthetic references indicates whether the RamiGlyph embedding is more consistent with continuous variation or discrete clustering. **E** Projection distributions and empirical cumulative distribution functions (ECDFs) for representative neighboring cluster pairs. The scaled dip statistic indicates the degree of separation between each pair, with larger values reflecting stronger evidence of discreteness. **F** PCA representation colored by the continuous morphology score. The score is defined by each cell’s normalized position along the fitted principal curve, ranging from simple branching (0) to complex branching (1). **G** Distribution of distances from individual cells to their projections on the principal curve, reflecting deviation from the main morphological trajectory. **H** Projection residuals across the continuous morphology score, reflecting deviation from the fitted principal curve. **I** Representative microglial morphologies selected along the continuous morphological trajectory. **J** Median standardized morphometric features across the morphology score, showing progressive changes in process extent and branching complexity. **K** Normalized morphology score distributions in held-out test cells from all CKp25 brain regions, the KXA experiment in S1, and CKp25 CB. Vertical lines indicate group medians. **L** Spearman correlations between the morphology score and classical morphometric features.

We then assessed whether weights trained on microglia could transfer directly to Neuron7. Without retraining, the transferred model reached 0.70-0.72 accuracy and 0.63-0.65 macro-F1 across K values, lower than the 0.78-0.82 accuracy and 0.73-0.78 macro-F1 obtained by training on neuronal data (Supplementary Fig.5D). Bootstrap UMAP and centroid-distance analyses likewise showed reduced but preserved separation among neuronal types (Supplementary Fig.5C). Confusion matrices showed the lowest classification accuracy for stellate and aspiny cells (Supplementary Fig.5E). Together with the results above, this indicates that RamiGlyph captures tree-structured morphology beyond microglia, and that pretrained weights offer a usable starting point for new cell types.

### A morphology score quantifies the continuum of microglial morphology

Having shown that RamiGlyph embeddings captured biologically meaningful morphology, we examined whether microglial morphology forms discrete subtypes or a continuum. Here, we used the consistent validation strategy as Weis et al. ^28^ In t-SNE space, microglial morphologies occupied a continuous landscape rather than forming clearly separated clusters (Fig. 5A). We tested this by fitting Gaussian mixture models in PCA space, calibrated against synthetic mixtures ranging from well separated to strongly overlapping components (Supplementary Fig.6A). If stable subtypes existed, clustering consistency should peak at the corresponding number of components. Across 100 repeated fits, however, the adjusted Rand index (ARI) declined monotonically with component number and showed no peak (0.96, 0.83, 0.69 and 0.44 at K = 2, 5, 10 and 80), whereas synthetic data with ten discrete components peaked clearly at K = 10 (ARI = 0.91; Fig. 5B, C). Density dips measure the valley depth between neighboring morphological clusters, with larger values indicating stronger separation (Fig. 5E; Supplementary Fig. 6B, C). The real data had a median within cluster spread to inter-centroid distance ratio of 0.32 (Fig. 5D). At this separation level, the maximum normalized density dip was only 0.11, below the 0.21 of matched synthetic data, indicating a substantial population of transitional cells (Fig. 5D). A maximum adjacency boundary dip of 0.129 suggested local enrichment, but the overall structure remained spectrum-like and continuous.

**Fig. 6.**
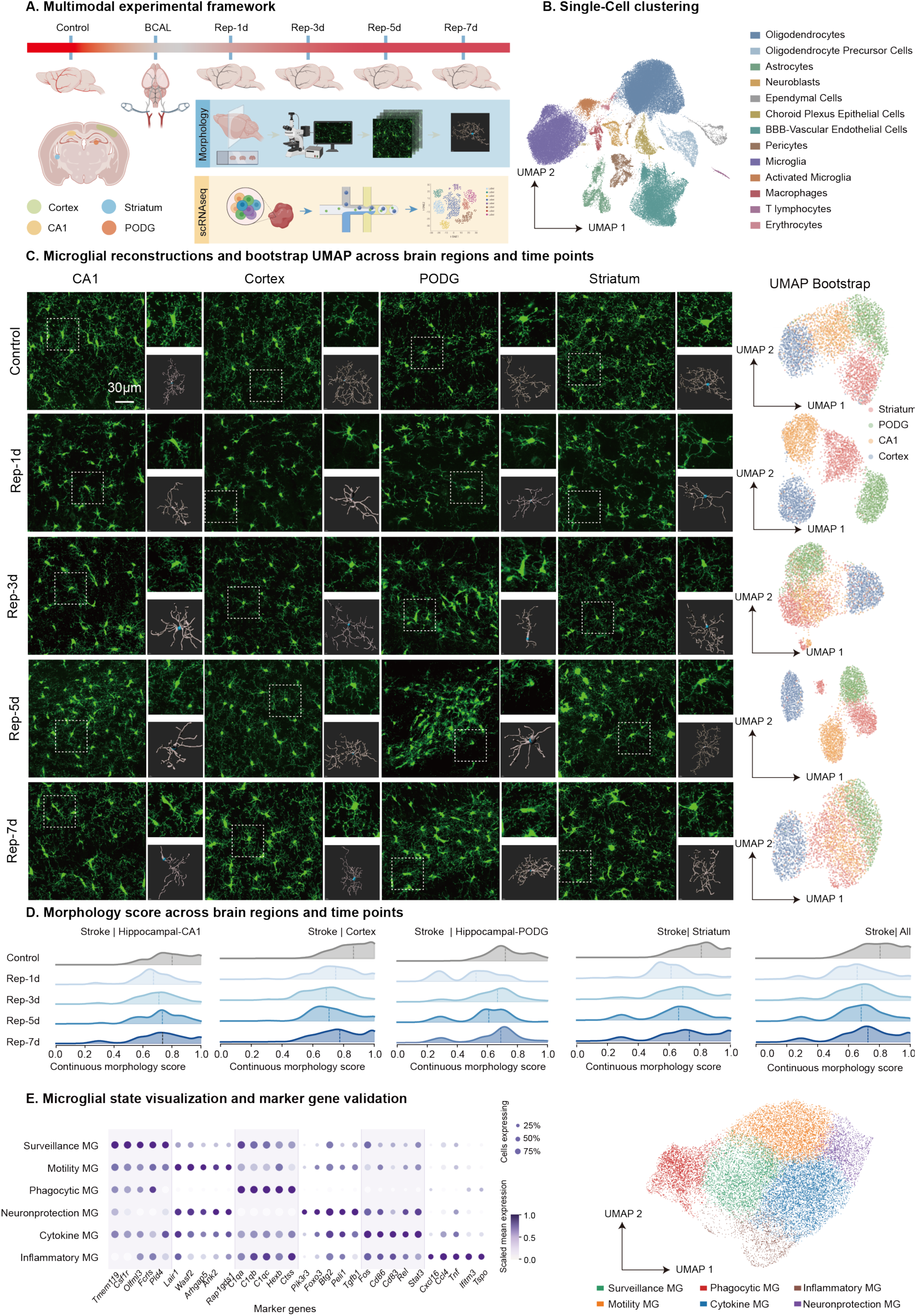
Spatiotemporal profiling of microglial morphology and transcriptional states after BCAL. **A** Experimental design of Control, BCAL and reperfusion stages (Rep-1d, Rep-3d, Rep-5d and Rep-7d), including four sampled brain regions and parallel morphological and single cell RNA sequencing analyses. **B** UMAP visualization of the integrated single cell transcriptomic dataset colored by major cell type annotations. **C** Representative fluorescence images and corresponding microglial reconstructions from hippocampal CA1, cortex, polymorphic layer of the dentate gyrus (PODG), and striatum across Control and reperfusion stages. Bootstrap UMAPs show regional organization of microglial morphology over time. **D** Morphology score distributions for each brain region and for all regions combined across the five stages. Scores range from simple branching (0) to complex branching (1). **E** Marker gene validation of major cell type annotations and visualization of microglial states. In the dot plot, dot size indicates the percentage of cells expressing each marker and color indicates scaled mean expression. Microglial UMAPs show six annotated states.

The morphological axis was constructed from a random subset of 5,000 training cells using the first 15 principal components (81.7% of variance). The resulting diffusion map formed a single connected component with no detectable branching. We therefore defined a continuous morphology score based on each cell’s normalized position along the principal curve. (0-1), oriented from simple to complex branching (Fig. 5F). The score varied smoothly across cells in PCA, t-SNE and UMAP space, and projection residuals were low (median 13.80, 95th percentile 25.25), indicating that the curve fitted most cells well (Fig. 5G, H; Supplementary Fig. 6D, E). Scores from 50 repeated subsamples were highly concordant with the reference fit (median Spearman’s ρ = 0.99), showing that the score robustly captures each cell’s position along the dominant morphological spectrum (Supplementary Fig. 6F).

To identify the morphological features underlying the continuous morphology score, we first examined representative cells sampled along the trajectory, which showed a gradual transition from relatively simple process to increasingly elaborate branching patterns as the score increased (Fig. 5I). A morphometric heat map revealed coordinated increases in process length, spatial extent, radial distance and branching related features along the same axis (Fig. 5J). We then quantified these relationships by permutation importance in a random forest regression model, which predicted scores accurately in held-out cells (R² = 0.95). Terminal point number was the strongest predictor (mean decrease in R² of 0.47), followed by convex hull surface area, total Sholl intersections, total length and peak Sholl intersections (Supplementary Fig. 6G, H). Correlation analysis identified a similar set of features, with the score correlating positively with total sholl intersections, total length and terminal-point number (Spearman’s ρ = 0.93-0.94) and negatively with Sholl decay slope (ρ = -0.49; Fig. 5L). The score therefore reflects coordinated variation in branching complexity, process length and spatial extent rather than a single morphometric feature.

We evaluated the continuous morphology score in 4,322 CKp25 and 345 KXA cells from the held-out test set. In CKp25, score distributions shifted progressively toward lower values from control to weeks 1, 2 and 6, in six of the seven brain regions, with CB as the exception (Fig. 5K; Supplementary Fig. 7A). In KXA, examined in S1, scores decreased with increasing dose and rebounded progressively during recovery (Fig. 5K). These changes were consistent between training and test sets, and the curve fitted test cells as well as training cells, so the shifts reflect genuine morphological change rather than fitting artefacts (Supplementary Fig. 7B, C). Together with the morphometric interpretation above, these results indicate that CKp25 progression and increasing KXA dose shifted microglia toward simpler branching. The rebound during recovery suggests that these changes may be reversible.

### RamiGlyph reveals microglial morphological trajectories from newly acquired ischemia reperfusion datasets

We next tested RamiGlyph on an independently acquired dataset from a different disease model. Global cerebral ischemia was induced in mice by bilateral common carotid artery ligation (BCAL), and brain tissue was collected at 1, 3, 5 and 7 days (Fig. 6A). Two-photon imaging and single-cell reconstruction yielded 5,690 microglia from four regions-cortex, striatum, hippocampal CA1 and the polymorphic layer of the dentate gyrus (PoDG)-for model evaluation (Fig. 6C).

We first assessed discriminative performance at the population and single-cell levels. Bootstrap analysis revealed that morphological separability among brain regions fluctuated over the course of reperfusion, peaking at 1 and 5 days and converging at 3 and 7 days (Fig. 6C). Centroid distance analysis showed that cortical microglia diverged most from the other regions at all time points except Rep-1d, with PoDG ranking highest at that stage (Supplementary Fig. 8A). Microglial morphology was distinguishable in all four brain regions at each of the five time points, and distances between the control group and each injury group exceeded those between any two injury groups (Supplementary Fig. 8B, C). At the single cell level, embeddings combined with morphometric features predicted pathological stage with accuracies of 0.62-0.72 using logistic regression and support vector machines (Supplementary Fig. 9A, B, C, D). RamiGlyph therefore generalizes to an independent dataset from a distinct disease model.

We next projected all cells onto the previously established reference trajectory. The trajectory covered the range of morphological variation in this dataset, with 93.87% of cells falling within the 99^th^-percentile residual threshold (Supplementary Fig. 9F). The median morphology score declined in all brain regions during early reperfusion, followed by recovery and fluctuation at later stages (Fig. 6D). Microglial branching complexity therefore follows a complex-simple-complex trajectory during ischemia reperfusion, confirming that the continuous morphological axis transfers across datasets and disease models (Supplementary Fig. 9E).

To further establish the coupling between microglial morphology and function, we generated a single cell transcriptomic dataset spanning control and 1, 3, 5 and 7 days after ischemia (Supplementary Fig. 10A). After quality control and integration, 75,901 cells were annotated into 13 cell types on the basis of canonical marker genes (Fig. 6B, Supplementary Fig. 10B, D). The 19,846 microglia recovered were further resolved into six transcriptional states, which formed a continuum rather than discrete subpopulations (Fig. 6E, Supplementary Fig. 10C). This atlas provides an independent molecular reference for linking morphology to function.

### Cross modal alignment predicts functional composition from microglial morphology after ischemia reperfusion

To determine whether microglial morphology encodes interpretable functional information, we developed a three stage framework comprising transcriptomic state definition, cross modal alignment and morphology-based functional prediction (Fig. 7A). We first built a biologically interpretable set of microglial functional pathways. Integrating GO and KEGG enrichment results, we selected 159 representative pathways and grouped them into eight functional states: antig-presenting, IFN-responsive, inflammatory, migratory-chemotactic, neuroprotective, phagolysosomal, proliferative and surveillance. Each state comprised 4-41 pathways and 60-441 genes (Fig. 7B). State-specific genes accounted for only 24.10%-78.70% of each state, indicating substantial sharing. Overlap was greatest between IFN-responsive and inflammatory states (Jaccard index 0.23), followed by inflammatory and neuroprotective (0.13) and migratory-chemotactic and neuroprotective (0.12). The eight states are therefore molecularly distinct yet not independent, and are better regarded as overlapping functional programs than as discrete subtypes (Fig. 7C).

**Fig. 7.**
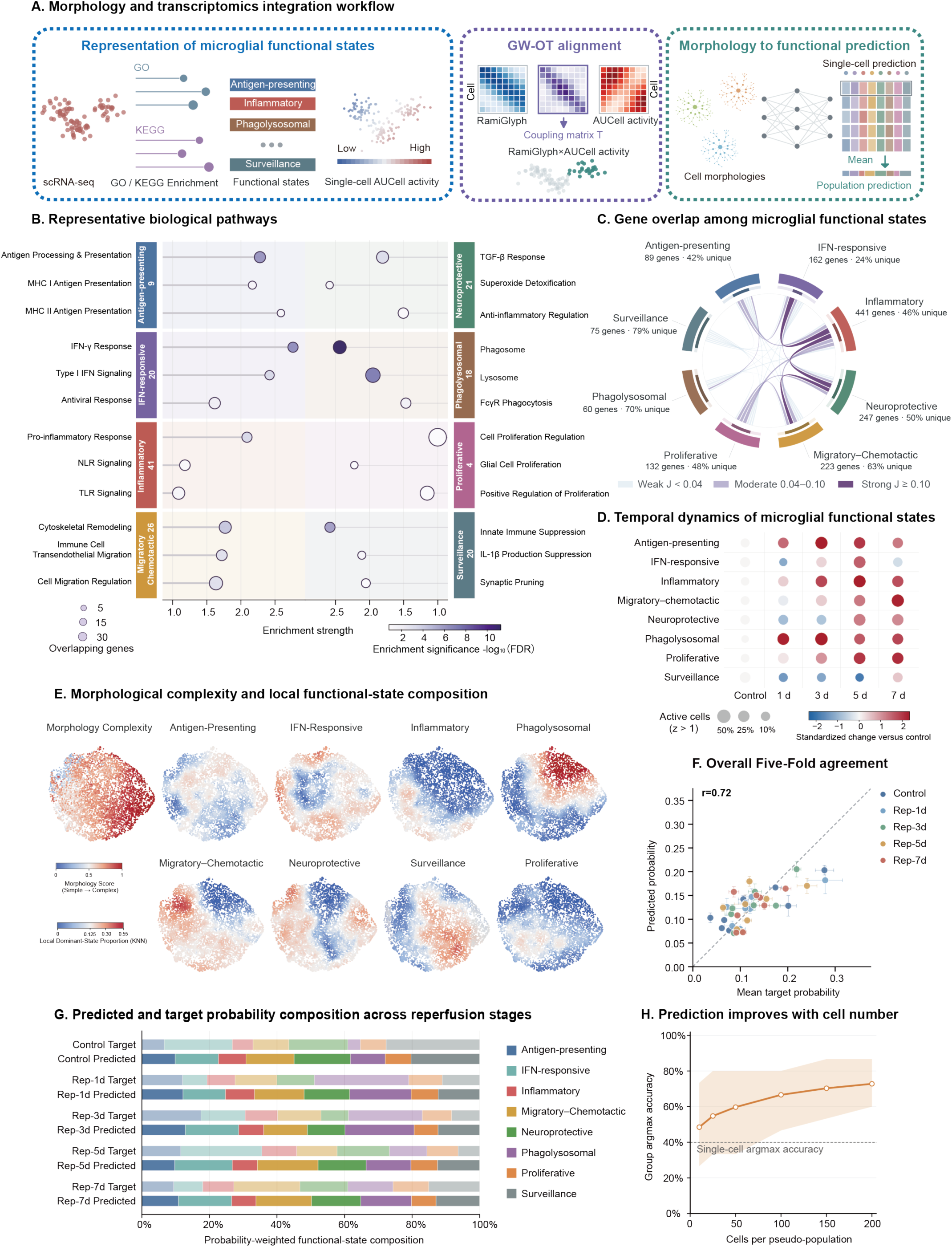
Morphology–transcriptomics integration predicts microglial functional states across reperfusion stages. **A** Workflow for integrating microglial morphology and transcriptomics. GO/KEGG enrichment defines transcriptomic functional states, AUCell quantifies their single cell activities, and GW-OT aligns RamiGlyph representations with AUCell activity through a coupling matrix. The aligned data are used for morphology - prediction at the single cell and population levels. **B** Representative pathways enriched across eight microglial functional states. Dot position indicates enrichment strength, dot size indicates the number of overlapping genes, and color indicates enrichment significance. **C** Chord diagram showing gene overlap among functional states. Labels indicate the total gene number and state specific fraction. Ribbon width indicates the number of shared genes, and ribbon color indicates Jaccard similarity. **D** Temporal changes in functional state activity across reperfusion stages. Dot size indicates the percentage of active cells, and color indicates the standardized change relative to Control. **E** RamiGlyph UMAP colored separately by the continuous morphology score and the local k nearest neighbor proportion of cells dominated by each functional state. **F** Correlation between predicted and target state probabilities across reperfusion stages in five fold cross validation. Error bars indicate standard deviations across folds, and Pearson’s indicates the overall correlation. **G** Target and predicted functional state compositions across reperfusion stages. **H** Bootstrap population level accuracy across different pseudo population sizes. Shading indicates the 95% bootstrap interval, and the dashed line indicates single cell accuracy.

We next applied Area Under the Curve cell score (AUCell) to score the eight functional states in each of the 19,846 microglia^29^. The states showed locally enriched yet overlapping distributions, with individual cells engaging multiple programs simultaneously (Supplementary Fig. 11A). These programs were then reorganized in a highly ordered temporal sequence. At 1 day, the phagolysosomal program was activated first, with active cells rising nearly eightfold (3.60% to 28.40%; Fig. 7D). Antigen-presenting activity increased in parallel (8.70% to 17.90%), whereas surveillance contracted (19.80% to 8.90%; Fig. 7D). This early response was amplified at 3 days, with marked activation of the inflammatory program. By 5 days, most programs were broadly activated and surveillance fell to its lowest level, marking the peak of the injury response (Fig. 7D). At 7 days, migratory-chemotactic and proliferative programs became dominant (33.30% and 24.20%), inflammatory and phagolysosomal activity remained elevated, and surveillance partially recovered (Fig. 7D). The microglial injury response is therefore not a single "activation" event. Instead, microglia undergo a functional succession, progressing from early clearance and antigen presentation, through broad immune activation, to late migration, proliferation and partial homeostatic recovery.

To align morphology with transcriptomic function in the absence of paired samples, we performed GW-OT independently within each condition (control and 1, 3, 5 and 7 days after reperfusion), mapping pathway activities onto the morphological embedding space (Fig. 7E; Supplementary Fig. 11C). In the representative 3-day condition, morphological distances correlated strongly with the transported transcriptomic distances (Spearman ρ = 0.892, n = 12,000 cell pairs; Supplementary Fig. 11B). Across time points, functional compositions inferred from morphology matched those measured from the transcriptome (Supplementary Fig. 11D). The alignment therefore preserved the internal structure of both spaces.

Finally, we tested whether the functional states inferred by GW-OT could be predicted from morphology alone. Using morphological embeddings of 5,690 cells as input and the GW-OT state probabilities as training targets, we trained a multi-task MLP and evaluated it by five-fold cross validation at the cell level (Fig. 7F; Supplementary Fig. 11E). Accuracy was 40.00% and macro-F1 33.80% (chance level 12.5%), indicating that morphology carries detectable functional information but is insufficient to assign the state of individual cells. Predictions became more accurate when probabilities were pooled across cells, rising from 48.5% for groups of 10 cells to 72.8% for groups of 200 (60.0%-86.7% across 500 resamplings at 200 cells; Fig. 7H). Predicted and target compositions agreed at every time point (Pearson r = 0.57-0.87; Fig. 7G; Supplementary Fig. 11F). Morphology therefore resolves functional state poorly in single cells but accurately at the population level, providing an indirect readout of population functional composition.

### The morphology function correspondence generalizes to amyloid pathology

To test whether the morphology function correspondence generalizes beyond ischemic injury, we applied the same framework to the 5×FAD model of amyloid pathology. UMAP and centroid distance analyses gave consistent results across the three stages in each brain region. In CB, CN and SN, the three stages occupied relatively distinct areas of the embedding. In FC and S1, controls were clearly separated from the two 5×FAD stages, whereas in DG and OB the stages overlapped substantially, indicating a more continuous morphological shift (Fig. 8A; Supplementary Fig. 12). Morphology was therefore distinguishable at the population level, though to differing degrees across regions.

**Fig. 8.**
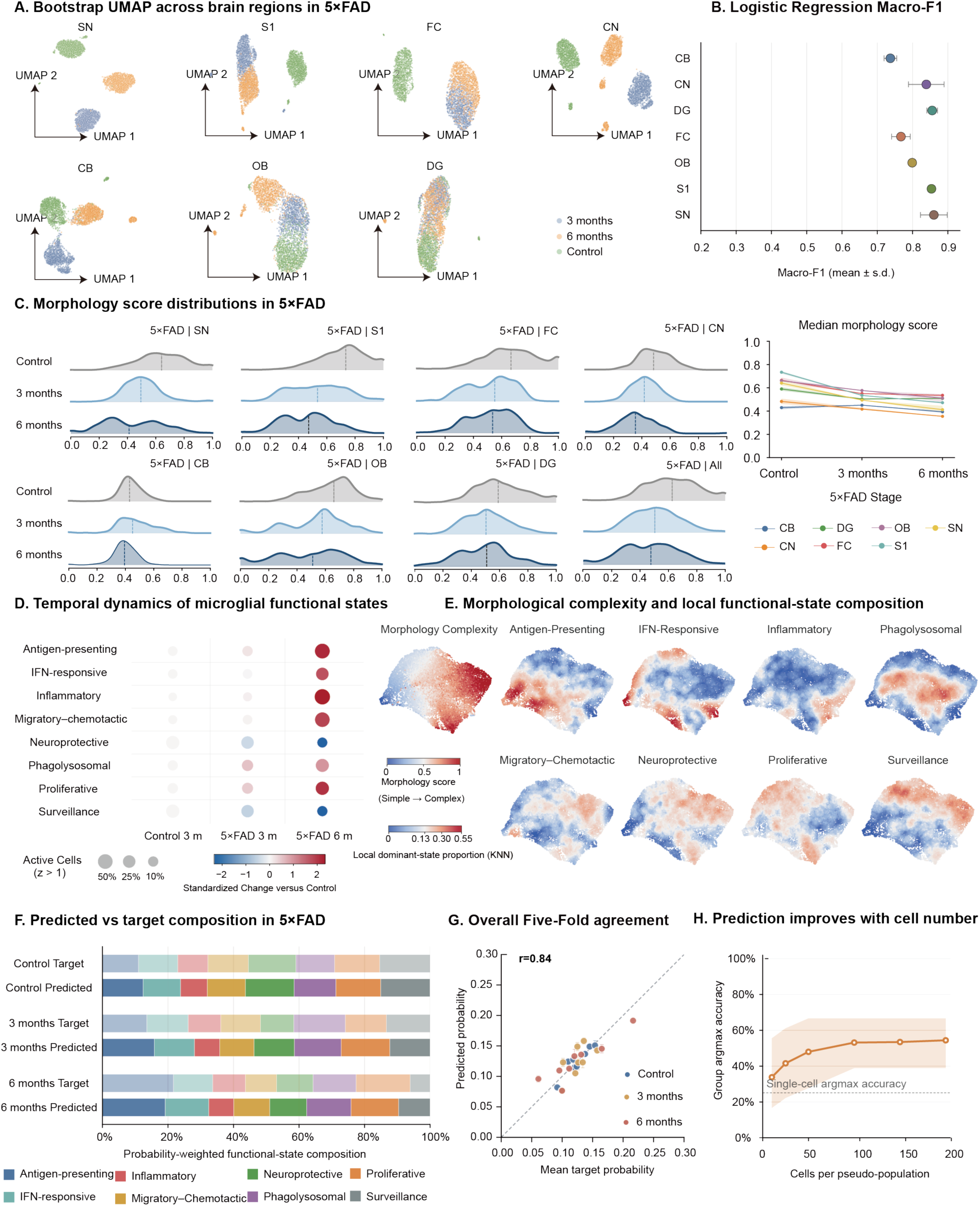
Regional morphology changes and functional state prediction in 5×FAD mice. **A** Bootstrap UMAPs of microglial morphology across seven brain regions (SN, S1, FC, CN, CB, OB and DG), colored by disease stage. **B** Disease stage classification from single cell morphology across brain regions using logistic regression. Points show mean macro F1 ± s.d. across five folds. **C** Fixed reference morphology score distributions and regional median trajectories across 5×FAD stages. Scores range from 0 (simple) to 1 (complex). Dashed lines indicate medians and shaded bands indicate 95% resampling intervals. **D** Temporal changes in eight functional state activities across Control, 5×FAD 3 and 6 month groups. Dot size indicates the percentage of active cells, and color indicates the standardized change from Control. **E** Continuous morphology score and local functional state composition mapped onto the morphology UMAP. Functional maps show the local k nearest neighbour proportion for each state. **F** Target and predicted functional state compositions across 5×FAD stages. **G** Correlation between predicted and target state probabilities across 5×FAD stages in five fold cross validation. Error bars indicate standard deviations across folds. **H** Bootstrap population level accuracy across pseudo population sizes. Shading shows the 95% bootstrap interval, and the dashed line shows single cell accuracy.

Projecting 15,593 microglia onto the reference morphological axis left only 1.61% with high trajectory residuals, indicating that the axis applies equally to the 5×FAD model (Supplementary Fig. 13C). Across all regions, the median morphology score declined from 0.64 in controls to 0.50 at 3 months and 0.48 at 6 months, indicating a progressive reduction in branching complexity with disease progression. Region-wise patterns were heterogeneous. CB showed little change and DG shifted early before plateauing, whereas the score declined monotonically in the remaining five regions (Fig. 8C).

Morphological embeddings were combined with morphometric features to predict disease stage in single cells. Performance was assessed by stratified five-fold cross validation within each region. Logistic regression reached 0.85-0.93 accuracy and 0.74-0.86 macro-F1 across the seven regions, with a linear SVM performing similarly (0.850-0.932 and 0.741-0.864, respectively; Supplementary Fig. 13A, B).

To relate morphology to function in the 5×FAD model, we integrated single cell transcriptomes from controls and 5×FAD mice at 3 and 6 months, identifying seven microglial transcriptional states (Supplementary Fig. 14A, B, C). AUCell scoring of the eight functional programs again produced locally enriched yet overlapping activity regions on the transcriptomic UMAP (Supplementary Fig. 15A). Phagolysosomal and proliferative programs were already elevated at 3 months. By 6 months, antigen-presenting, IFN-responsive, inflammatory, migratory-chemotactic and proliferative programs were further upregulated, whereas surveillance and neuroprotective programs continued to decline (Fig. 8D). Microglial function therefore shifted from homeostatic surveillance and neuroprotection toward immune activation, phagocytosis and proliferation over the course of 5×FAD pathology.

We then applied GW-OT within each disease stage to align the morphological and transcriptomic data. The eight functional programs formed partly overlapping activity regions in the morphological UMAP, with KNN analysis confirming enrichment of several states within specific morphological neighborhoods (Fig. 8E; Supplementary Fig. 15C, D). Alignment quality was supported by the strong correlation between morphological and transported transcriptomic distances at 3 months (Spearman ρ = 0.867, n = 12,000 cell pairs; Supplementary Fig. 15B).

Finally, we tested whether the eight state probabilities defined by GW-OT could be predicted from morphology alone. Predicted and target compositions correlated at r = 0.79-0.85 across five folds (pooled r = 0.84), and at 0.92, 0.66 and 0.91 for control, 3 months and 6 months (Fig. 8F, G; Supplementary Fig. 15E, F). Prediction of discrete single-cell states was more limited (accuracy 0.25, macro-F1 0.21; chance level 0.13). In randomly sampled cell groups, accuracy for the dominant state rose from 0.34 to 0.54 as group size increased from 10 to 200 cells (Fig. 8H). Microglial morphology and functional state therefore show a quantifiable correspondence that is more robust at the population scale than for discrete single-cell labels.

## Discussion

We developed RamiGlyph, a self-supervised model that directly learns representations of 3D microglial morphology. RamiGlyph embeddings capture biologically meaningful variation across conditions and generalize across disease datasets and neuronal morphologies. The embedding space revealed a microglial morphological continuum, enabling a unified score for cross dataset comparison. Building on this morphological representation, we used GW-OT to align the morphology space with transcriptomic states. This alignment further enabled morphology-based inference of functional states, revealing greater predictive reliability at the population level.

RamiGlyph captures rich phenotypic information at both the single cell and population levels. This capability is mainly attributed to the topology branch, which learns global process organization beyond predefined topological descriptors^10^. In addition, a Transformer-based direction aware attention mechanism incorporates soma relative orientation of branching points to capture spatial morphological features^30^. Together, these features are particularly suited to microglia, whose processes rapidly extend and retract in response to environmental perturbations^1,31,32^. RamiGlyph uses a SwAV-based strategy that learns separate prototypes in the topology and structure branches and aligns prototype assignments across views^24^. This process groups morphologically similar cells around shared prototypes, capturing the continuous heterogeneity of microglial morphology.

Earlier studies often classified microglia into categories such as ramified, rod-like, activated, and amoeboid, whereas more recent approaches have emphasized continuous morphological variation^8,33,34^. For example, MorphOMICs revealed brain region dependent morphological spectra^10^, while MorphoCellSorter ranked cells along feature-based morphological gradients^35^. Our analysis further suggested that local morphological enrichment does not necessarily define discrete cluster boundaries, as validated by synthetic data. RamiGlyph therefore defines a continuous morphology score along its major variation axis, enabling quantification of branching changes without relying on predefined morphometric features.

Our results indicate that microglial morphology reflects functional state more reliably at the population level. Pathway enrichment identified eight biologically meaningful programs consistent with previous transcriptomic studies ^6,34^. GW-OT linked these programs to morphology by aligning relational structures across unmatched cells^20^ . This strategy is conceptually related to GW-based cross modal integration methods such as SCOT and Pamona ^36,37^ . Although functional prediction was limited for individual cells, performance improved substantially after population-level aggregation. Thus, morphology appears to encode functional state primarily as a population level signal rather than a precise single cell readout.

RamiGlyph showed strong generalization across independent disease datasets and cell types. Although RamiGlyph was trained without condition or brain-region labels, its embeddings consistently distinguished these biological contexts across multiple classifiers and resampling analyses. This discriminative structure was preserved in independent BCAL and 5×FAD datasets, indicating that the learned representation was not specific to the training dataset. Notably, RamiGlyph also achieved performance comparable to SGTMorph trained directly on Neuron7^15^, despite the framework being originally designed for microglia. Together, these findings suggest that RamiGlyph captures transferable morphological representations across disease contexts and tree-like cellular morphologies.

Functional state analysis further revealed distinct trajectories between acute and chronic disease contexts. After BCAL, early morphological simplification coincided with increased phagolysosomal and inflammatory programs, whereas later structural recovery was accompanied by increased surveillance and neuroprotective programs. This temporal diversification is consistent with single cell studies showing heterogeneous inflammatory, phagocytic, and repair associated microglial states after ischaemic injury^38,39^. In 5×FAD mice, phagolysosomal programs were enriched early, followed by increased antigen presenting and proliferative states with disease progression. Similar transitions from homeostatic towards disease associated, MHC II expressing, and proliferative microglial states have been reported during amyloid accumulation^40–42^. Importantly, the contrast between BCAL and 5×FAD indicates that similar morphological simplification can accompany distinct functional programs depending on disease context and stage.

Several limitations should be considered. First, the current evaluation primarily reflects cell level representation, as cells from the same animal were not treated as an independent grouping factor. Second, transcriptomic data were obtained by dissociation, limiting fine spatial information within brain regions for morphology transcriptome alignment. Third, GW-OT establishes probabilistic rather than direct cell to cell correspondence, which may introduce uncertainty into morphology-based functional state inference. Future studies integrating 3D morphology with spatial or paired molecular profiling will be needed to directly validate these morphology function relationships.

RamiGlyph provides a generalizable framework for learning transferable morphological representations across tree-like cells and for quantifying changes in morphology along a continuous axis. In ischemia reperfusion and amyloid pathology models, RamiGlyph further showed that morphology can predict disease stage and infer functional states at the population level. Building on these results, RamiGlyph may provide a basis for morphology-based drug screening and functional state prediction in these disease models. In new disease datasets, pretrained RamiGlyph representations could be aligned with unmatched data from other modalities to enable phenotype based functional prediction.

## Methods Datasets

SWC reconstructions of 23,637 microglia were obtained from the Siegert laboratory archive on NeuroMorpho.Org, spanning adult homeostasis, repeated KXA anesthesia and recovery, ovariectomy and CK-p25 induced neurodegeneration, and were split 80:20 within each condition into training and evaluation sets^10^. The Neuron7 dataset (1,393 reconstructions, seven neuronal classes) was provided by the MorphoGNN authors^14^. Transcriptomic data for the 5×FAD analyses were obtained from GEO (GSE296768, GSE195518, GSE229373 and GSE273690). Morphological and transcriptomic data from the BCAL model were generated in this study. Reconstructions from 5×FAD mice, Neuron7 and the BCAL dataset were all held out from training and used for validation.

## RamiGlyph architecture

### Graph construction

Each SWC reconstruction was converted into a soma-rooted attributed graph, with sampling points as nodes and parent child relationships as bidirectional edges. Each node carried a nine-dimensional attribute vector (three-dimensional coordinates, radius and one-hot node type), with coordinates normalized to [0,1] and Euclidean distances between adjacent nodes stored as edge lengths. Three soma-referenced descriptors were added per node: soma distance (number of edges from the soma), branch level (number of bifurcations along that path) and primary branch identity (which soma-adjacent subtree the node belongs to), which together allowed the direction-aware attention module to distinguish centripetal, centrifugal and inter-branch node pairs. Reconstructions exceeding 1,000 nodes were simplified by removing nodes of degree below three in random order, protecting the soma and its immediate neighbors, so that all branch points and graph connectivity were retained.

### Dual-view augmentation

Two augmented views were generated from each graph for self-supervised training. Terminal branches were truncated at randomly selected positions, with all distal nodes removed. Branch points were preserved, leaving the branching structure intact. This procedure was repeated up to 20 times per view or until 500 nodes remained, followed by subsampling to 500 nodes. Coordinates were perturbed by mirroring the cell along a randomly chosen axis with probability 0.5 and adding independent jitter drawn uniformly from ±0.5 μm. Perturbations were applied to the original coordinates, after which coordinates were normalized and the soma-referenced descriptors recalculated on the augmented graph.

### Topo-branch encoding

The topological branch encodes the overall branching organization from the persistence image of the same augmented graph. Because microglial processes extend outward from the soma, we used the shortest-path edge distance from the soma as the filtration value, so that persistence reflects the order in which branches emerge. Zero-dimensional extended persistent homology gives a persistence diagram *D*_0_ = {(*b_n_*, *d_n_*)}, where *b_n_* and *d_n_* are the birth and death values of the *n*-th connected component. Each pair was mapped onto a 100 × 100 grid with a Gaussian kernel of bandwidth 1, weighted by its persistence |*d_n_* − *b_n_*|, and normalized to [0,1]. The resulting single-channel image was encoded by four convolutional blocks (3 × 3 convolution, batch normalization, ReLU and pooling; 32, 64, 128 and 256 channels), the first three with 2 × 2 max pooling and the fourth with adaptive average pooling, yielding a 256-dimensional representation after flattening. Parameters were shared between views.

### Structural branch encoding

The structural branch characterizes morphology from three complementary aspects: local connectivity, 3D geometry and soma-referenced branch hierarchy. Each node *i* is represented by ℎ*_i_*, obtained by concatenating its nine-dimensional attributes projected to 224 dimensions with its random-walk positional encoding projected to 32 dimensions. Edge labels were projected to 256-dimensional embeddings e_ij_.

The encoder stacks four identical blocks, each passing the same node representation through three parallel pathways. The local pathway propagates information along SWC-defined edges:

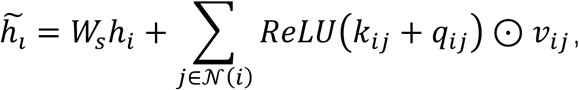

Where *k_i_*_j_, *q_i_*_j_ and *v_i_*_j_ are projected from the concatenated endpoint representations and edge embedding, and the ReLU term modulates each neighbor’s contribution^22^. Because connectivity indicates which nodes are linked but not their spatial arrangement, the geometric pathway adds an E(n)-equivariant network whose edge messages incorporate squared Euclidean distances, while node coordinates are iteratively updated across successive EGNN layers^23^. Both pathways are confined to graph edges, whereas the direction-aware attention pathway allows interactions between all node pairs.

Its content score 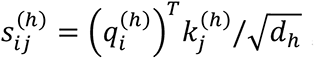, (four heads, *d_h_* = 64) ignores relative position within the process. We therefore classified node pairs by branch level *l* and primary branch identity *b* into four directional relations: centripetal (*b_i_* = *b_j_*, *l_i_* > *l_j_*), centrifugal (*b_i_* = *b_j_*, *l_i_* < *l_j_*), intra-branch (*l_i_* = *l_j_*, *b_i_* = *b_j_*) and inter-branch (*b_i_* ≠ *b_j_*). Each relation carries a learnable embedding *E_r_*_,ℎ_ and head specific weight *ω_r_*_,ℎ_, indexed by the level difference *δ_i__j_* = |*l_i_* − *l*_j_|; and a second bias encodes the soma distance difference, giving

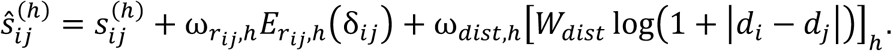

Because all node pairs are evaluated, complexity scales quadratically with node number, so augmented graphs were limited to 500 nodes. The three pathway outputs are concatenated and combined by a gating network (768 → 256 → 3) whose softmax weights allow their relative contributions vary across nodes, followed by a residual connection and feed-forward network. After four blocks, node representations are mean pooled and projected to the 256-dimensional cell-level structural representation.

### Self-supervised prototype learning

Both encoders were trained with a branch-specific prototype-learning objective adapted from SwAV^24^, encouraging the two augmented views of a cell to yield consistent prototype assignments without labels. Each branch had its own set of 30 learnable prototypes, frozen for the first 200 steps. For branch *r* and view *m*, the *L*_2_ -normalized encoder output 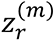 was multiplied by the unit-norm prototype matrix *Cr* to give scores 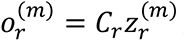, which the Sinkhorn–Knopp algorithm converted into balanced soft assignments 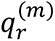 (entropy regularization 0.05, three iterations). Assignments were treated as fixed targets, and the assignment from one view supervised the prototype prediction of the other:

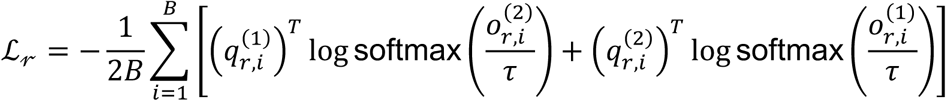

where *B* is the minibatch size and *τ* = 0.1 . The two branch losses were weighted equally and no feature queue was used.

The model contains 7.46 million trainable parameters and was trained for 3,000 optimization steps using AdamW (weight decay, 1 × 10^−5^) with a batch size of 100. The structural and topological branches used maximum learning rates of 3 × 10^−4^ and 5 × 10^−3^, respectively. Learning rates were linearly increased during the first 1,000 steps and subsequently reduced by cosine decay to 1% of their maximum values. Mixed-precision training and gradient clipping at a global norm of 1.0 were applied, with all random seeds fixed at 42. Training comprised approximately 16 effective epochs and 600,000 augmented graph views on an NVIDIA GeForce RTX 4090 GPU.

### Baseline methods

All comparison methods were retrained on our data rather than used as released models. SGTMorph, MorphoGNN, Tree-MoCo, MorphVAE, MorphOMICs and conventional morphometric features were evaluated on the same partition (18,913 training and 4,724 held-out cells), with no labels used during representation learning^10,12–15^. Each method retained its native preprocessing, training objective and author-recommended hyperparameters. For RamiGlyph, the structural and topological representations were concatenated to 512 dimensions; in the fusion analysis the morphometric block was scaled by 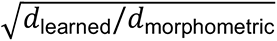 before concatenation so that both blocks contributed comparably to distance calculations.

### K nearest neighbor evaluation

Each representation was evaluated with a k nearest neighbor (KNN) classifier, using the training partition as reference samples and the held-out validation partition for evaluation only. Labels were assigned by unweighted majority voting among the k nearest training samples (k = 1, 3, 5 and 10). Learned representations are *L*_2_ -normalized and were compared by cosine distance, whereas MorphOMICs and morphometric features carry physical units and were standardized on training statistics before Euclidean comparison. Accuracy and macro-F1 were used as performance measures.

### Ablation analysis

Ablation variants were trained separately, each with one component removed and all other settings unchanged: the topological branch encoder, direction-aware attention, the equivariant graph neural network or the graph convolutional network for architectural ablations; branch pruning, spatial reflection, coordinate jitter, rotation or translation for augmentation ablations. All variants were evaluated under the same KNN protocol (k = 5). For component *m* and task *t*, the performance decrease was 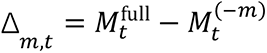, where *M* is macro-F1, and was averaged across tasks. Each component’s relative contribution was its mean decrease divided by the summed mean decreases within its group; these percentages describe how the total decrease is distributed, not the size of the decrease itself.

### Bootstrap resampling and visualization

To assess the stability of group-level morphology, cells were pooled and stratified by experimental condition. Because cell numbers varied across conditions, pseudo-bulk populations of fixed size were constructed by sampling 15 cells with replacement and retaining their mean dimensional representation; this was repeated 1,000 times per condition. Replicates were projected onto the first 50 principal components and visualized by t-SNE (perplexity 50) and UMAP (50 neighbors, cosine distance, min-dist = 1.0, spread = 3.0)^43,44^. All quantitative comparisons were performed in the original 512-dimensional space, with the two-dimensional projections used only for visualization.

### Centroid distances and multivariate separation analysis

Condition centroids were calculated from bootstrap representations in the dimensional embedding space, and pairwise Euclidean distances assembled into a distance matrix visualized by average-linkage hierarchical clustering. Separately, single-cell Euclidean distances were partitioned into within- and between-group pairs; separation was summarized by the difference in mean distance and quantified by Cliff’s delta, with a one-sided Mann-Whitney U test comparing the two distributions^45^. Multivariate separation was assessed by PERMANOVA with 999 permutations, reporting pseudo-F, R² and permutation P values, and PERMDISP with the same number of permutations was used to test for differences in within-group dispersion^26,46^.

### Single-cell classification (LR and SVM)

Predictive information in the single-cell representations was evaluated by stratified fivefold cross-validation using multinomial logistic regression (LR; *L*_2_, *C* = 1, up to 1,000 iterations) and a linear support vector machine (SVM; one-vs-rest, *L*_2_, squared-hinge loss, *C* = 1, primal formulation, up to 5,000 iterations), fitted on identical splits directly on the representations without PCA projection^27^. Each cell was predicted only when held out, giving out-of-fold predictions across the full dataset. Accuracy and macro-F1 are reported as mean and standard deviation across folds, with per-class performance and row-normalized confusion matrices computed from these predictions.

### Discrete versus continuous structure

Because no threshold defines "discrete" a priori, we calibrated against synthetic mixtures of known separation, following the strategy of Weis et al^28^. A ten-component diagonal-covariance Gaussian mixture model (GMM) was fitted to the training embeddings in 50-dimensional PCA space (covariance regularization 10^−4^, 200 EM iterations, five initializations); its component means *μ_k_* and weights *pi_k_* generated synthetic data as *x_i_* = *μ_zi+_* *λd_c_*_ℎ*ar*_*ε_i_*, where *z_i_* ∼ *Categorical*(*π*), *ε_i_* ∼ *N*(0, *I*) and *d_c_*_ℎ*ar*_ is the mean nearest-centroid distance. Nine continuity levels (*λ*= 0.05–1.00) were examined with three datasets each, matched to the real data in size. Cluster stability was the mean pairwise adjusted Rand index across 100 spherical-covariance GMM fits with different initializations, trained on up to 6,000 cells and evaluated on up to 5,000 held-out cells at *K* = 2, 5, 10, 20, 40, 60 and 80. Local separation was quantified by Hartigan’s dip statistic along centroid-connecting axes and by the spread-to-separation ratio 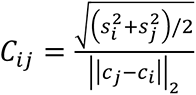 between nearest-neighbor components. Measured values were positioned against the synthetic series; no fixed threshold was applied.

### Continuous morphology trajectory and score

Trajectory inference used the first 50 principal components. An adaptive-kernel diffusion map (up to 5,000 training cells, 30 nearest neighbors, α = 0.5, t = 1) initialized the cell ordering, after which a 300-point principal curve was fitted iteratively. All cells were projected onto this fixed curve and scored by normalized arc-length position, 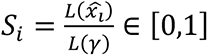, with residuals 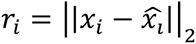 above the 99th training percentile flagged. Direction was set by branching-complexity features so that 0 denotes simpler and 1 more complex branching. Robustness was checked across 50 subsamples and at 15, 30 and 50 nearest neighbors. For application to other datasets the trajectory was frozen: scores were obtained by projection alone, without refitting, reorientation or renormalization, keeping values comparable across datasets.

### Morphological correlates of the score

To determine which morphological properties the score captures, we calculated Spearman correlations with NeuroM derived morphometric features (Benjamini-Hochberg adjusted)^47^. Trends were visualized by binning cells along the score axis and taking the median standardized feature value per bin. We then tested how well these features predict the score, using random-forest regression with fivefold cross-validation and permutation importance in held-out cells. Segment-level measurements were excluded, as they scale with SWC sampling density rather than with morphology. Representative cells with below-median trajectory residuals were selected at evenly spaced positions along the score axis. A second set spanned the 5^th^-95th percentiles of convex hull surface area, peak Sholl intersections and total arbor length.

### Bilateral common carotid artery ligation

Global cerebral ischemia was induced in adult Cx3cr1^GFP/+^ mice (18-25 g, 2-4 months) by bilateral common carotid artery ligation (BCAL) as previously described^48^. Mice were anesthetized with ketamine (150 mg/kg) and xylazine (25 mg/kg) i.p. The bilateral common carotid arteries were then exposed and ligated with 6-0 sutures for 60 min before release for reperfusion. Occlusion was verified by laser speckle contrast imaging (RFLSI III, RWD Life Science), with a ≥75% reduction in cerebral blood flow within 10 min defining successful ischemia. Control mice received the same anesthesia. All procedures were approved by the Ethics Committee of Experimental Animals, Jiaxing University (approval NO. JUMC2025-161).

### Two-photon imaging and three-dimensional reconstruction

Mice were anesthetized as above and transcardially perfused with ice-cold PBS followed by 4% PFA, and brains were sectioned at 60 μm. Cx3cr1^GFP/+^ microglia were imaged on a two-photon microscope (Olympus FV1000, 25×/1.05 NA water-immersion objective) as z-stacks at 0.75 μm intervals across 40 optical planes, in the cortex, striatum, hippocampal CA1 and PoDG; every fifth section was sampled. Imaging was performed on 25 mice, five each for control and 1, 3, 5 and 7 days after reperfusion.

Stacks were imported into Imaris 10.2.0 (Bitplane) and microglial processes traced in three dimensions with the filament-tracing module. New starting points were detected with a largest diameter of 12 μm and seeding points of 1 μm, and disconnected segments removed with a filtering smoothness of 0.6 μm. Cells lying at the image border and therefore only partially traced were removed manually. Traced filaments were exported in .ims format and converted to SWC using xyz2swc (https://neuromorpho.org/xyz2swc/ui/)^49^.

### Single-cell RNA sequencing

Separate cohorts were used for sequencing. Mice were anesthetized as above and transcardially perfused with ice-cold PBS, and the cortex, striatum and hippocampus dissected in the same buffer. Fifteen mice were used in total, with tissue from three mice pooled per time point (control and 1, 3, 5 and 7 days after reperfusion). Samples were stored in MACS Tissue Storage Solution (Miltenyi Biotec) before enzymatic dissociation into single-cell suspensions, with viability exceeding 85% in all sample.

Libraries were prepared using the DNBelab C Series Single-Cell Library Prep Set (MGI, #1000021082). Droplets were generated from the single-cell suspension, followed by emulsion breakage, bead collection, reverse transcription and cDNA amplification to generate barcoded libraries. Indexed libraries were constructed according to the manufacturer’s protocol, quantified using a Qubit ssDNA Assay Kit (Thermo Fisher Scientific, Q10212) and sequenced on a DNBSEQ platform (BGI, Shanghai).

### Single-cell RNA-seq processing and annotation

Analyses were performed in Scanpy (v.1.11.5)^50^, with the BCAL and 5×FAD datasets processed independently through the same workflow. Cells with fewer than 200 genes or more than 5% mitochondrial transcripts were removed, together with sample-specific thresholds on gene and transcript counts. Counts were normalized to 10,000 per cell, log-transformed, and the 2,000 most variable genes selected before PCA and Harmony batch correction^51^.

Neighborhood graphs (15-20 neighbors) were used for UMAP and Leiden clustering (resolution 0.2-0.4). Cell types were assigned manually from lineage markers identified by Wilcoxon rank-sum tests; microglia were then isolated and the workflow repeated for subclustering. Low-quality clusters and putative doublets were excluded during annotation.

### Construction of microglial functional gene sets

Differentially expressed genes were identified for each microglial subtype against all others (one-versus-rest, adjusted *P* < 0.05, |log₂FC| > 0.25), followed by over-representation analysis (ORA) using GO Biological Process 2025 and KEGG 2019 Mouse gene sets^52^. Enriched pathways were manually grouped into eight microglial functional states according to their annotated biological functions and published microglial literature. For each pathway, gene symbols were mapped to mouse orthologues, restricted to genes detected in at least 0.5% of microglia, and filtered to 5-50 genes per pathway with housekeeping, ribosomal and mitochondrial genes excluded. The resulting gene sets were stored as GMT files and used for both the BCAL and 5×FAD analyses.

### AUCell-based functional activity scoring

Functional activity was quantified per cell using AUCell implemented in pySCENIC^29^. The complete expression matrix was used as input, with equal gene weights within each pathway and AUC normalization disabled. Pathway-level scores were averaged across pathways assigned to the same functional state, yielding an 8-dimensional activity profile for each cell. For UMAP visualization, activity values were standardized separately within each state using robust z-scores. For temporal summaries, conventional z-scores were calculated across cells within each state, and cells with z > 1 were classified as active. Mean AUCell activity and active-cell fractions were summarized for each condition.

### Gromov-Wasserstein alignment

Morphological embeddings (512 dimensions) and AUCell state activities were aligned by entropically regularized Gromov-Wasserstein optimal transport, performed independently within each matched condition^20^. Within-domain dissimilarities were cosine distance for morphology and Euclidean distance for pathway activities, symmetrized and scaled to [0,1], with uniform marginals. Optimization used the proximal-point solver in the Python Optimal Transport package (ε = 5 × 10⁻³, 1,000 iterations, tolerance 10⁻⁸), retaining the best of five initializations. Transcriptomic activities were transferred to morphological cells by barycentric projection, the highest standardized activity taken as each cell’s dominant state. Fidelity was assessed by Spearman correlation of pairwise distances before and after transport across 12,000 cell pairs. The coupling represents a probabilistic correspondence between populations, not a one-to-one matching of cells.

### Morphology-based prediction of functional states

A multilayer perceptron with three residual SwiGLU blocks was trained to predict the eight GW-imputed functional states from the 512-dimensional morphology embedding alone^53^. Supervision combined the dominant state as a hard label with soft targets derived from the fold-standardized state activities, under equally weighted cross-entropy and Kullback-Leibler losses. Models were optimized with AdamW (learning rate 2 × 10⁻⁴, batch size 512) and evaluated by fivefold cell level cross validation stratified by condition and dominant state, with all predictions out-of-fold. Population compositions were obtained by averaging out-of-fold probabilities within each group, and pseudo-populations of 10-200 cells were sampled 500 times.

### Code availability

The source code for RamiGlyph, together with scripts for model training, morphology complexity score and downstream analyses, is available at https://github.com/quy21/RamiGlyph. Analyses were performed in Python 3.9 using standard open-source packages, including GUDHI, NumPy, pandas, scikit-learn, SciPy, PyTorch, PyTorch Geometric and umap-learn.

## Acknowledgements

This research was supported by the Zhejiang Provincial Natural Science Foundation of China under Grant No. LQN26H090013 and the International Scientific Research Cooperation Seed Fund of the International Campus, Zhejiang University, for the project entitled “Establishment and Application of a Multimodal Deep Learning Model for 3D Liver Organoids”. The authors thank all members of the laboratory for their support.

We would like to thank Make Zhao, Jingjing Zhao, Jiahua Peng, and Rao Fu, students at Zhejiang University, as well as Hao Gao (Northwest Minzu University), for their valuable technical support and helpful discussions.

## Author contributions

Y. Qu, Y. Chi and J. R. Liu conceived the project. Y. Chi and J. R. Liu supervised the study. Y. Qu designed the model architecture and conducted downstream functional analyses. Y. Qu and T. Lan performed the scRNA-seq analyses. J. R. Liu performed the BCAL experiments and image acquisition. J. Z. Xu performed cell extraction. Y. M. Xiao, J. L. Liu and Q. T. Qian contributed to code implementation and debugging. All authors provided feedback on the manuscript and reviewed the final version.

---

**Supplementary Fig. 1.**
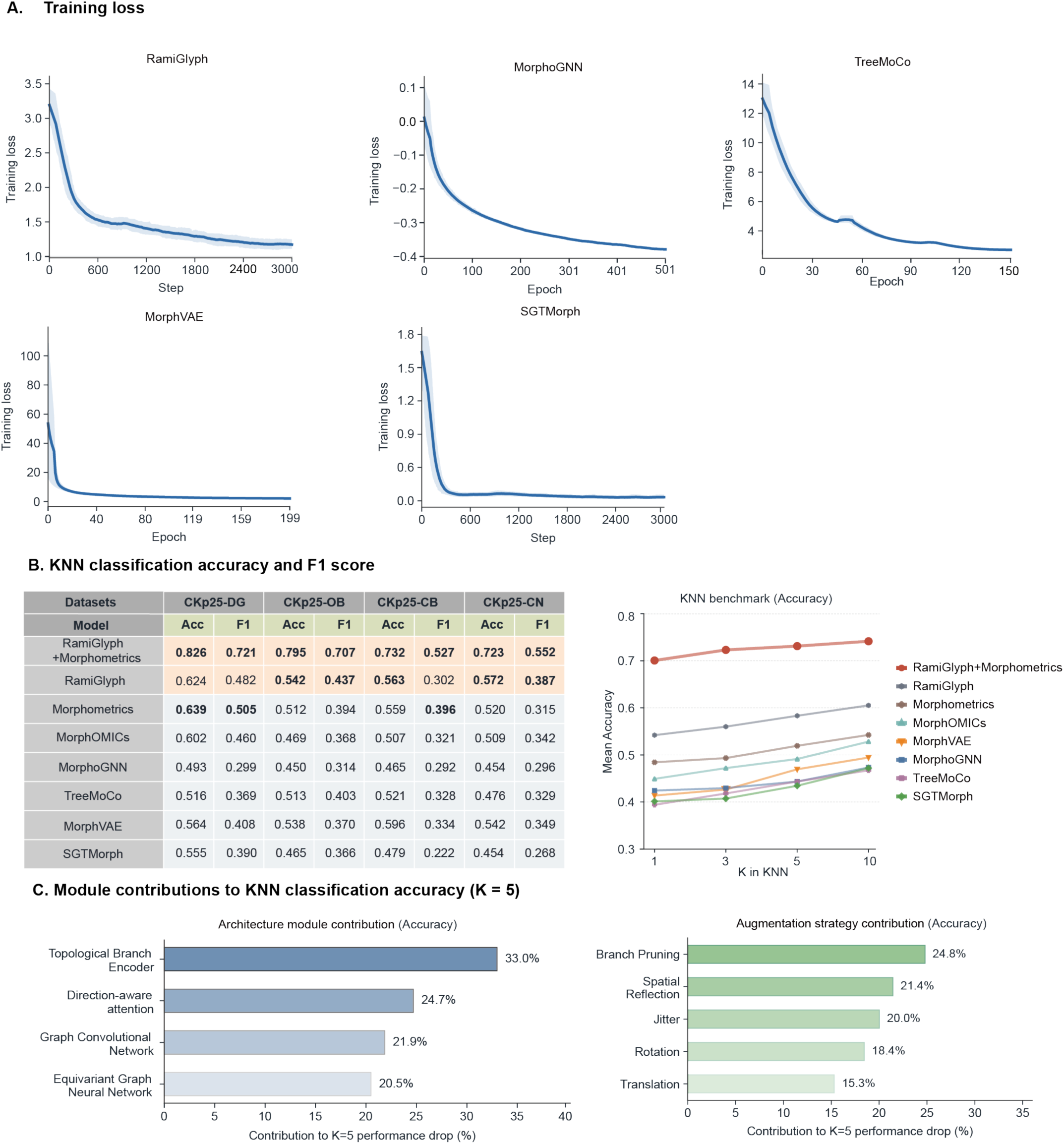
Training convergence, KNN benchmarking and module contributions. **A** Training loss trajectories for RamiGlyph and four representative deep learning models. **B** KNN classification performance at K = 5 for CB, CN, DG and OB, reported as accuracy (ACC) and macro F1; bold values indicate the best performing model. The line plot shows mean accuracy across ten evaluation datasets for K = 1, 3, 5 and 10. **C** Ablation analysis of module contributions to KNN accuracy at K = 5. Bars show the mean decrease in accuracy after removing each architectural module or augmentation strategy relative to the full model, averaged across datasets.

**Supplementary Fig. 2.**
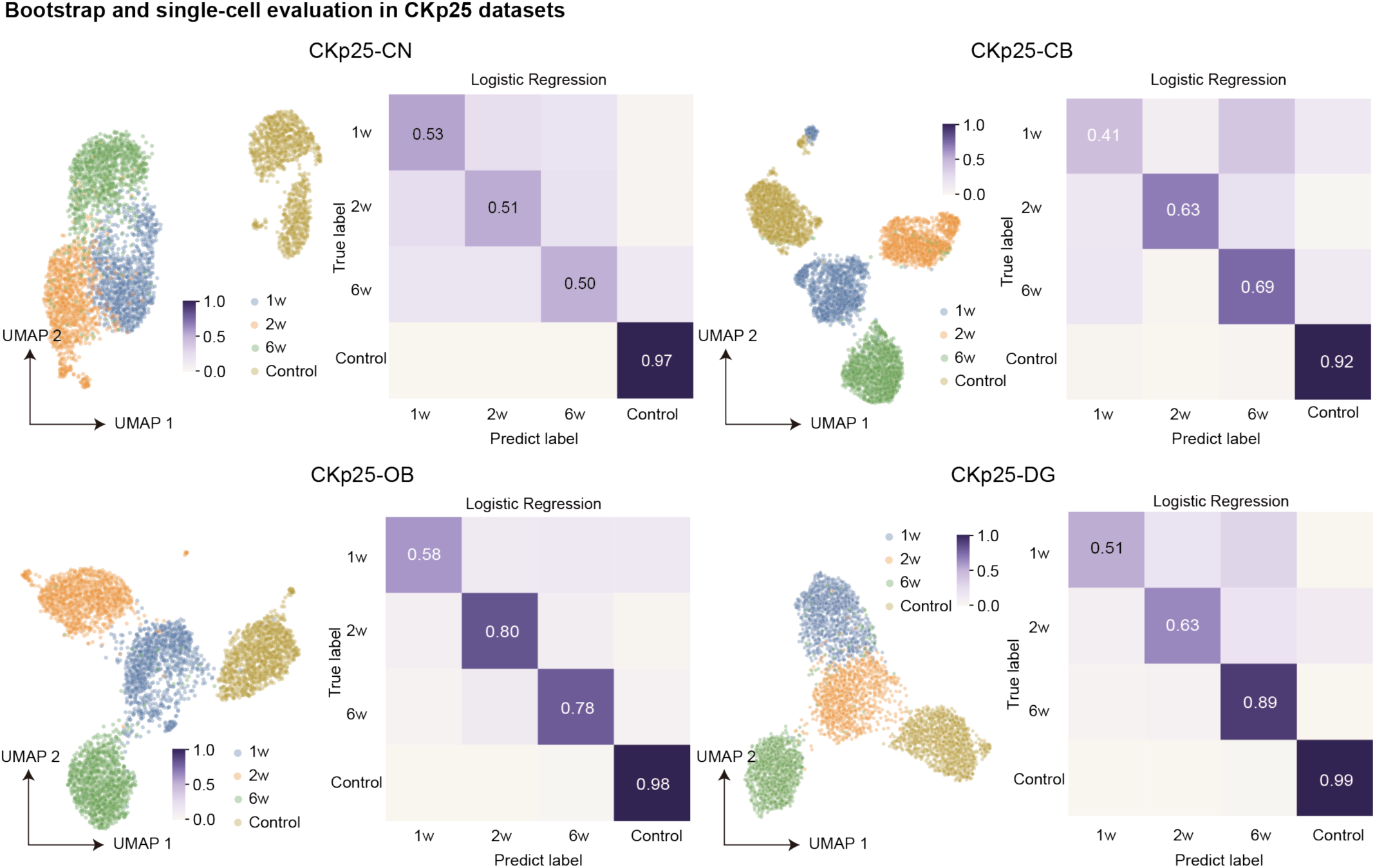
Bootstrap and single cell evaluation of CKp25 datasets. UMAP projections of bootstrap aggregated embeddings and logistic regression confusion matrices for the CN, CB, OB and DG regions. For each group, 1,000 bootstrap embeddings were generated by resampling 15 cells with replacement. Logistic regression predictions were obtained using five fold cross validation. Numbers on the diagonal indicate classification accuracy.

**Supplementary Fig. 3.**
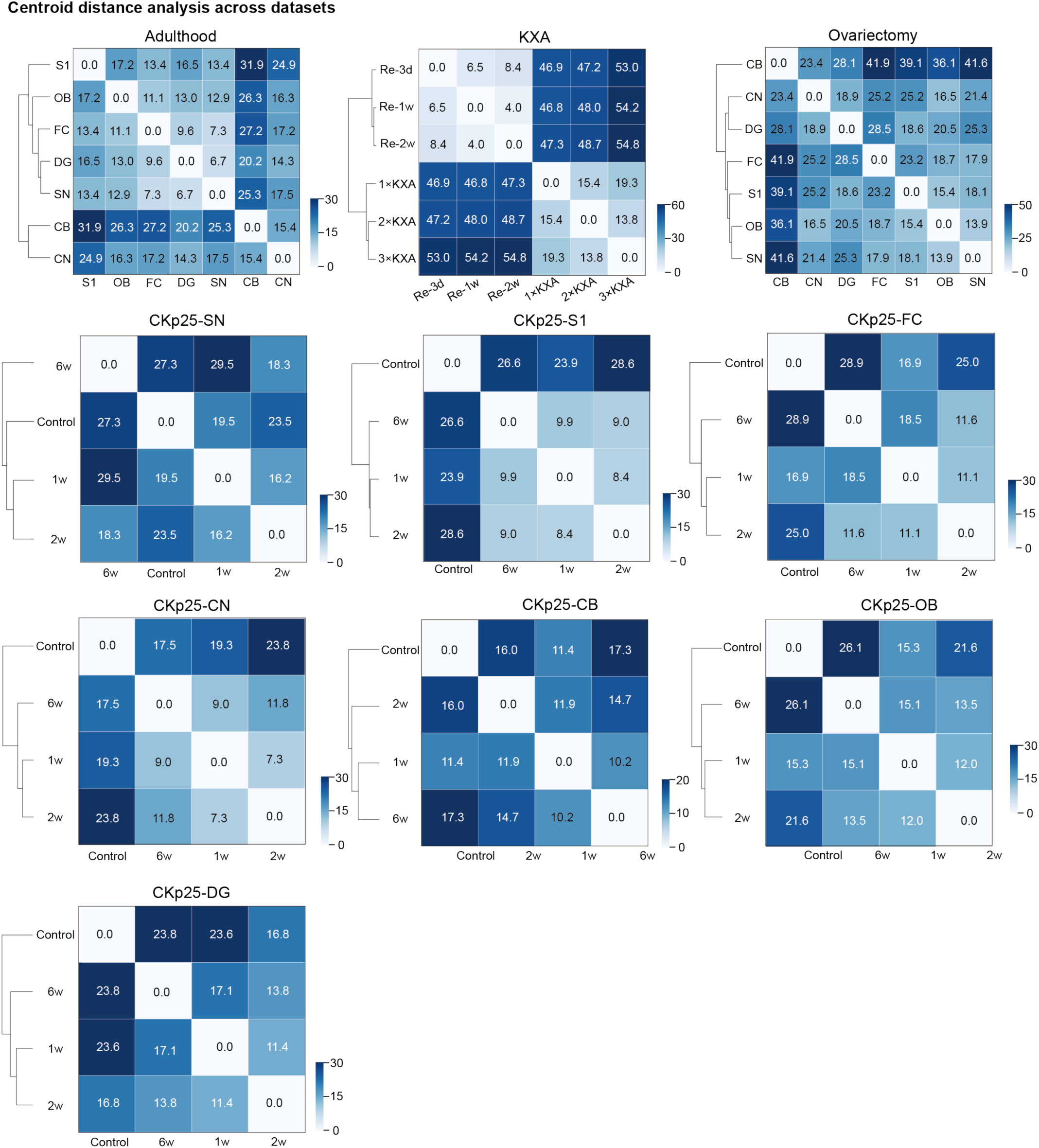
Centroid distance analysis across datasets. Heatmaps show pairwise Euclidean distances between bootstrap centroids for the Adulthood, KXA and Ovariectomy datasets and seven CKp25 brain regions. Dendrograms show hierarchical relationships among groups. Cell values indicate centroid distances, with darker blue representing greater morphological separation

**Supplementary Fig. 4.**
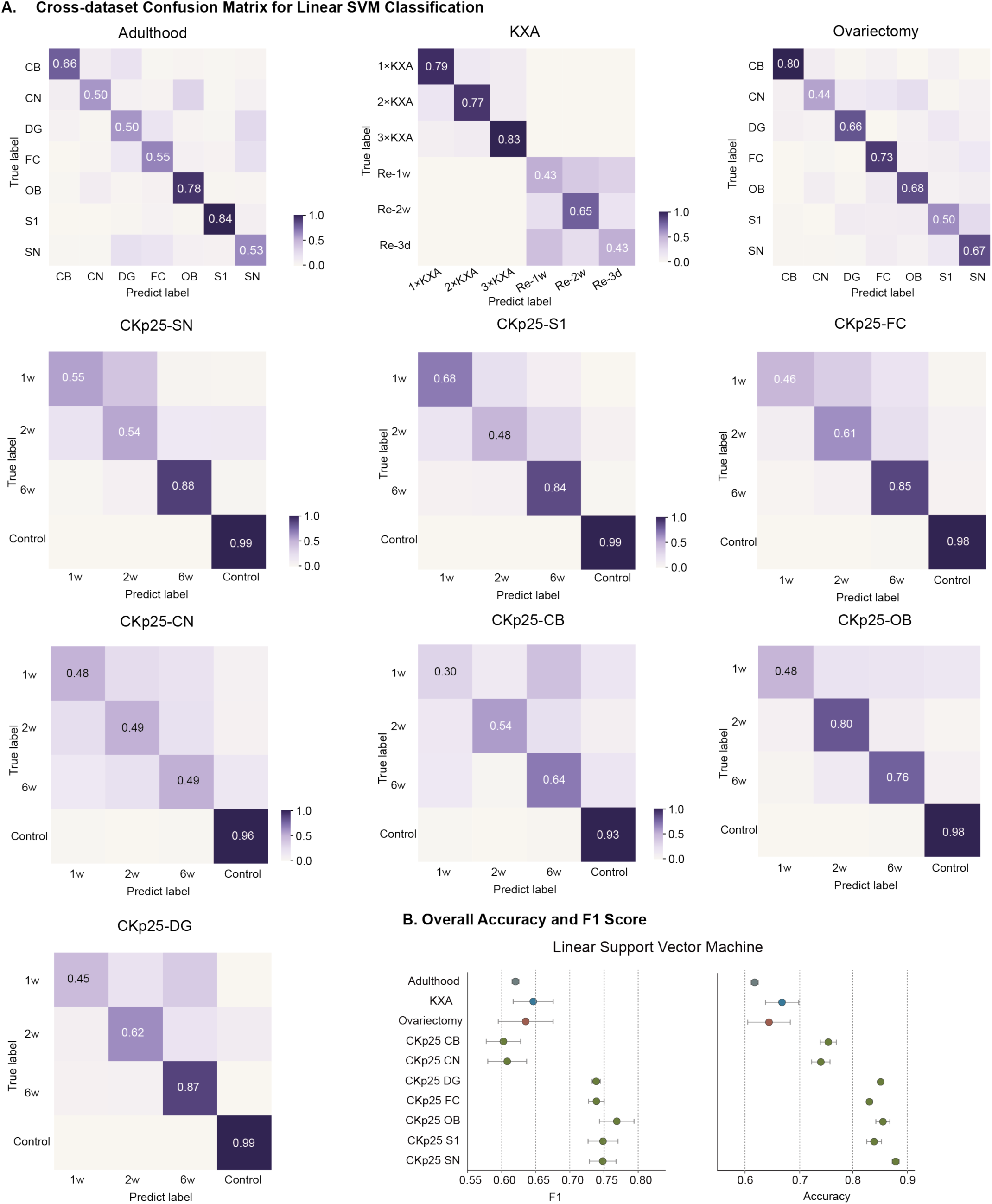
Single cell classification performance of linear support vector machines. **A** Row normalized confusion matrices from five-fold stratified cross validation for the Adulthood, KXA and Ovariectomy datasets and seven CKp25 brain regions. Numbers on the diagonal indicate classification accuracy. **B** Macro F1 score and overall accuracy across datasets. Points and error bars represent the mean ± s.d. over five folds.

**Supplementary Fig. 5.**
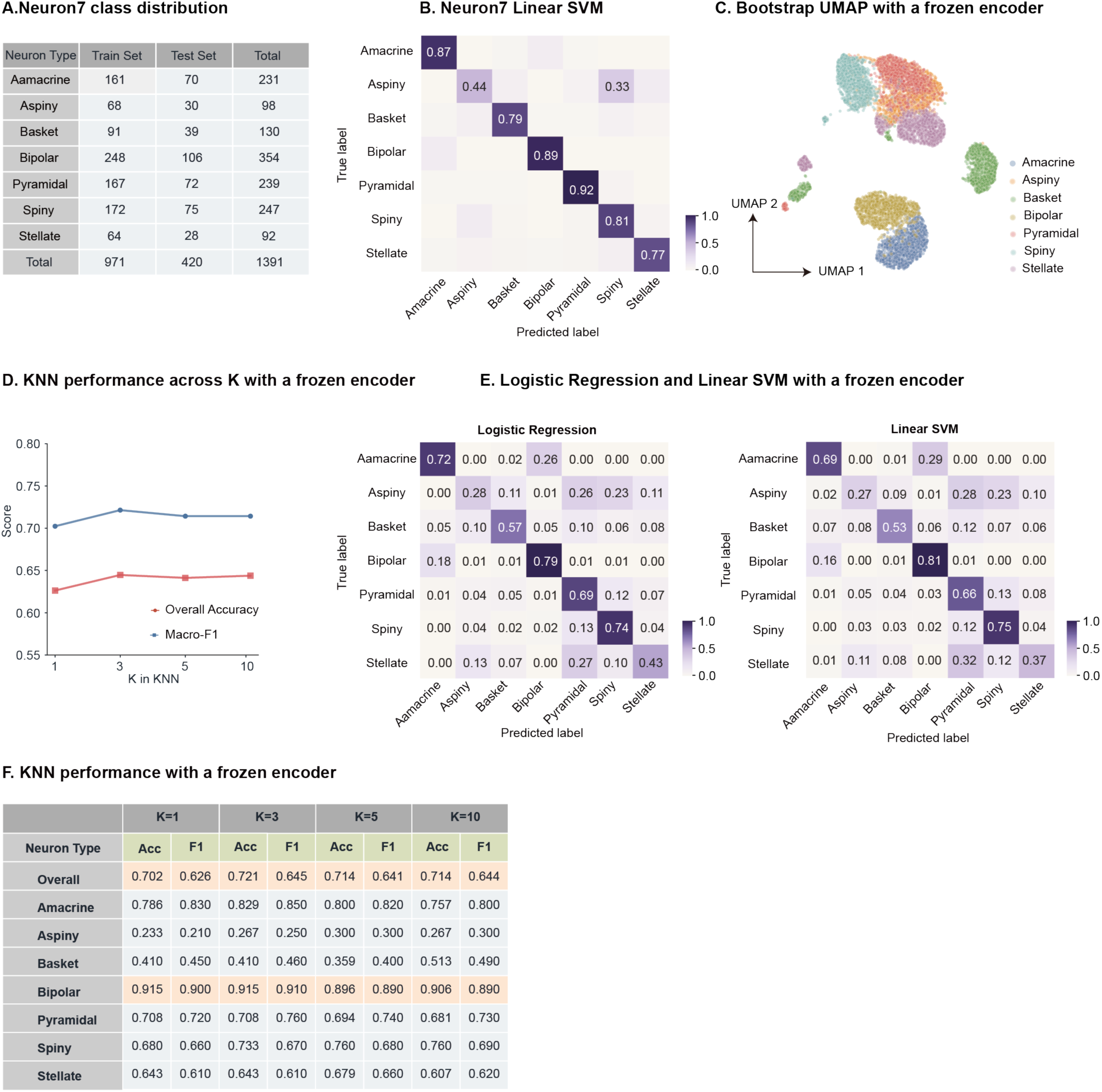
Dataset composition and frozen encoder evaluation on Neuron7. **A** Distribution of the seven neuronal classes across the training and test sets. **B** Confusion matrix from five-fold stratified cross validated linear SVM predictions using embeddings learned with Neuron7 specific training. **C** UMAP projection of bootstrap aggregated embeddings extracted using the frozen RamiGlyph encoder pretrained on microglial datasets. For each class, 1,000 bootstrap embeddings were generated by resampling 15 neurons with replacement. **D** Overall accuracy and macro F1 of KNN classification using embeddings from the frozen encoder for K = 1, 3, 5 and 10. **E** Confusion matrices from five-fold cross validated multinomial logistic regression and linear SVM using embeddings extracted by the frozen RamiGlyph encoder. **F** Overall and class specific KNN accuracy (ACC) and macro F1 using the frozen encoder across different K values.

**Supplementary Fig. 6.**
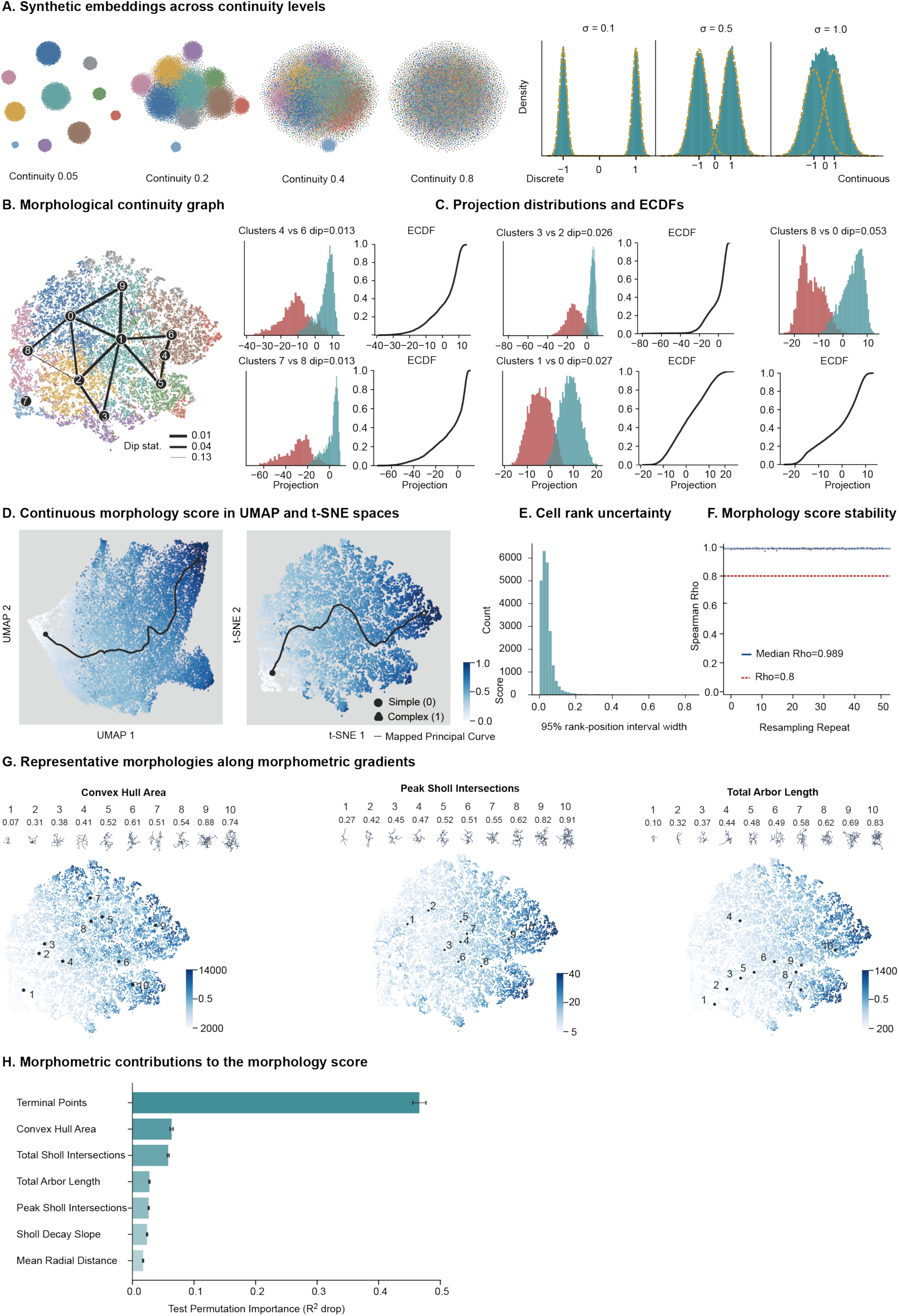
Validation of morphological continuity and the continuous morphology score. **A** Synthetic reference embeddings generated at continuity levels of 0.05, 0.2, 0.4 and 0.8, together with one-dimensional synthetic distributions at σ = 0.1, 0.5 and 1.0 illustrating increasing overlap between components. Teal histograms show the combined distributions, and orange dashed lines indicate the individual component densities. **B** Morphological continuity graph displayed on the clustered embedding. Numbered nodes indicate clusters, and edges connect neighbouring clusters. Edge thickness represents the corresponding dip statistic, with thicker edges indicating stronger separation. **C** Pairwise projection distributions and corresponding ECDFs for representative neighbouring cluster pairs. Shown are clusters 4 versus 6 (dip = 0.013), 7 versus 8 (dip = 0.013), 3 versus 2 (dip = 0.026), 1 versus 0 (dip = 0.027), and 8 versus 0 (dip = 0.053). **D** Continuous morphology score mapped in UMAP and t-SNE spaces. Black curves indicate the mapped principal curve, with circles and triangles marking the simple (0) and complex (1) ends, respectively. **E** Distribution of per cell 95% rank position interval widths, reflecting uncertainty in individual cell positions along the continuous morphology trajectory. **F** Stability of the continuous morphology score across 50 resampling repeats, quantified by Spearman correlation with the reference score. The blue line indicates the median correlation (ρ = 0.989), and the red line indicates ρ = 0.8. **G** Embedding distributions for convex hull area, peak Sholl intersections and total arbor length, respectively, with ten representative morphologies selected along each feature gradient. **H** Random forest permutation importance of seven morphometric features for predicting the continuous morphology score in the test set (*R*² = 0.95), quantified by the decrease in test *R*² after feature permutation.

**Supplementary Fig. 7.**
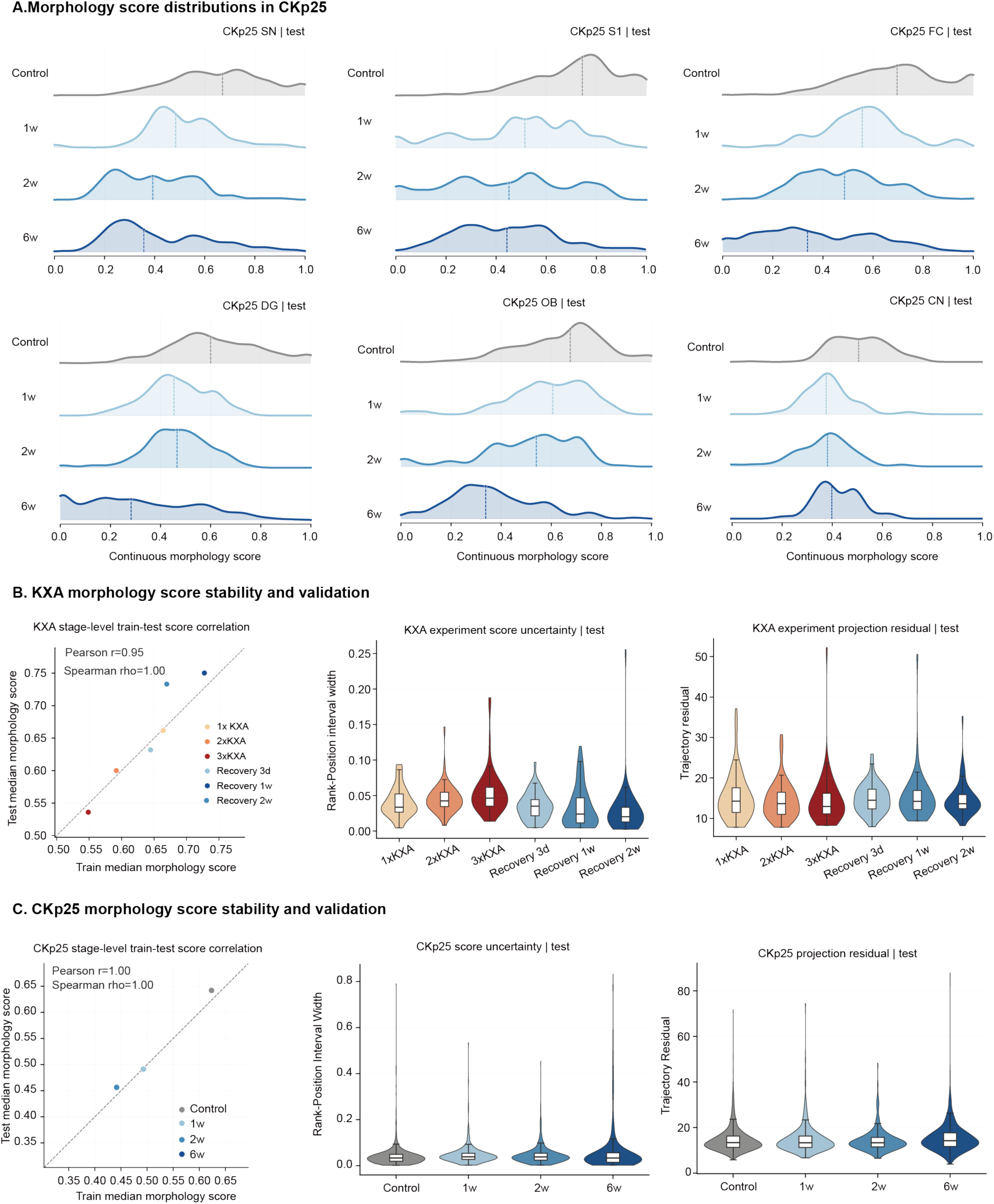
Regional morphology score distributions and test set validation in CKp25 and KXA. **A** Normalized morphology score distributions in CKp25 test cells across six brain regions at Control, 1 week, 2 weeks and 6 weeks. Vertical lines indicate group medians. **B** Validation of KXA morphology score stability. Stage level median morphology scores are compared between the training and test sets. Per cell rank uncertainty is quantified by the width of the 95% rank position interval, and principal curve projection residuals reflect the deviation of individual cells from the fitted morphological trajectory across KXA and recovery stages. **C** Validation of CKp25 morphology score stability. Stage level median morphology scores are compared between the training and test sets. Per cell rank uncertainty is quantified by the width of the 95% rank position interval, and principal curve projection residuals reflect the deviation of individual cells from the fitted morphological trajectory across CKp25 stages.

**Supplementary Fig. 8.**
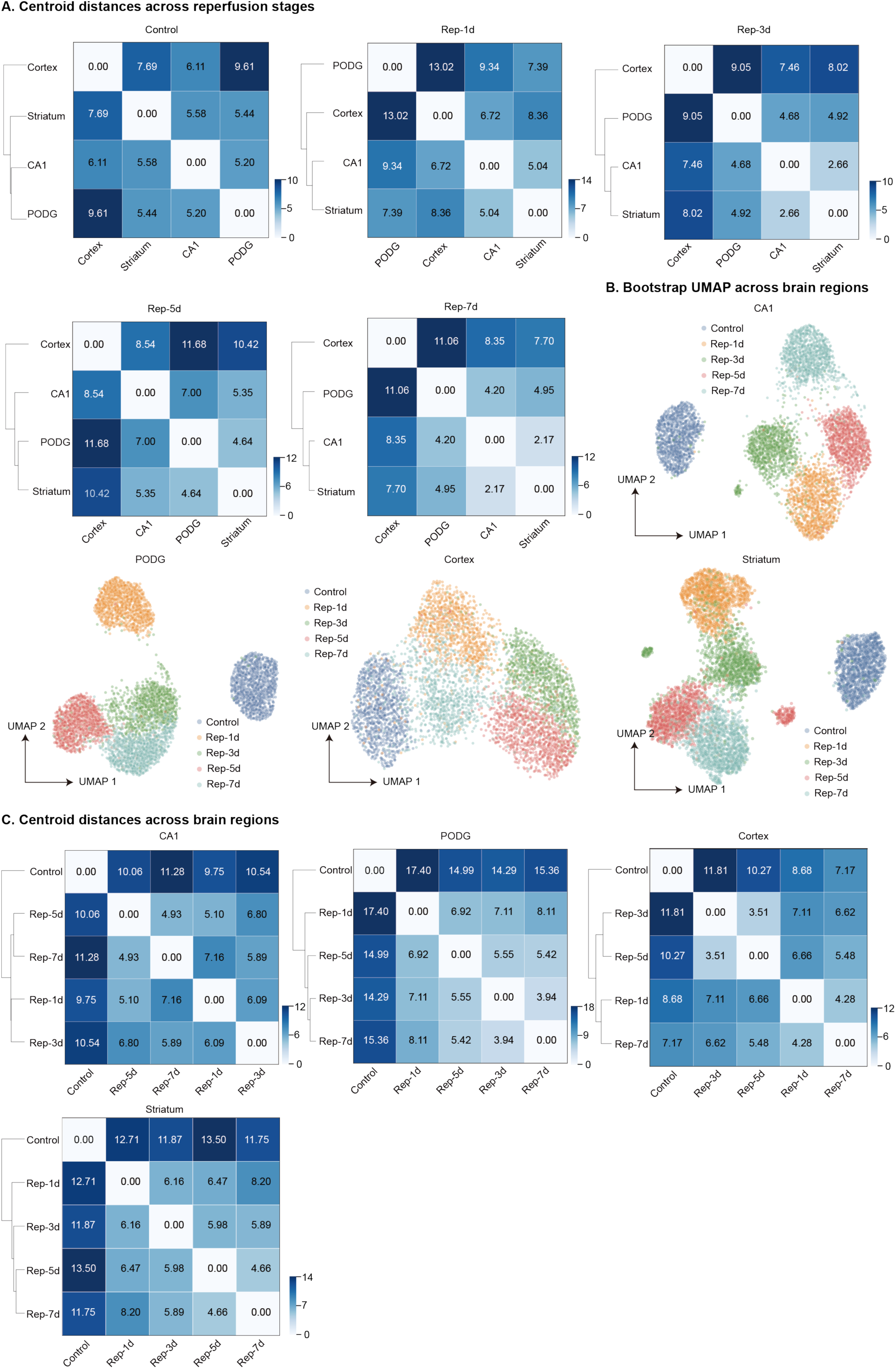
Centroid distances and bootstrap UMAPs of microglial morphology across BCAL stages. **A** Pairwise Euclidean distances between regional centroids at Control, Rep-1d, Rep-3d, Rep-5d and Rep-7d. Dendrograms show average linkage hierarchical clustering. **B** Bootstrap UMAPs for CA1, PODG, cortex and striatum, colored by reperfusion stage. **C** Pairwise Euclidean distances between stage centroids within each brain region.

**Supplementary Fig. 9.**
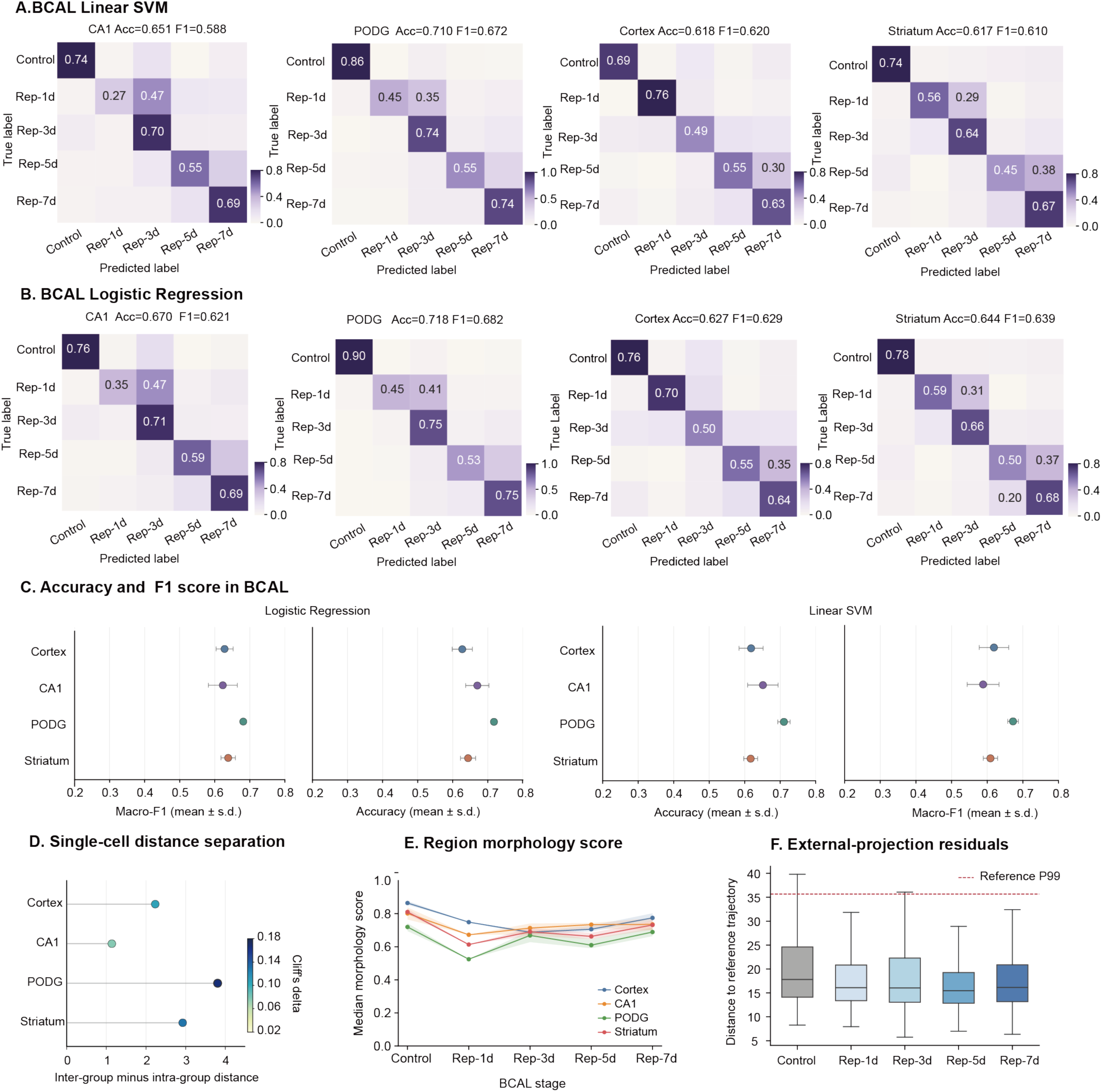
Regional separability and fixed reference projection of microglial morphology across BCAL stages. **A** Confusion matrices for linear support vector machine classification of Control, Rep-1d, Rep-3d, Rep-5d and Rep-7d cells within CA1, PODG, cortex and striatum. **B** Corresponding confusion matrices for multinomial logistic regression classification. **C** Regional classification accuracy and macro F1 for logistic regression and linear support vector machine models. **D** Single cell distance separation across brain regions, calculated as the mean inter stage distance minus the mean intra stage distance. Point color indicates Cliff’s delta. **E** Median fixed reference morphology scores across BCAL stages in cortex, CA1, PODG and striatum. Shaded bands indicate 95% intervals obtained by cell balanced resampling. Scores range from simple branching (0) to complex branching (1). **F** Distances of externally projected cells from the fixed reference trajectory across BCAL stages. Boxes show medians and interquartile ranges, and the red dashed line indicates the reference P99 residual threshold.

**Supplementary Fig. 10.**
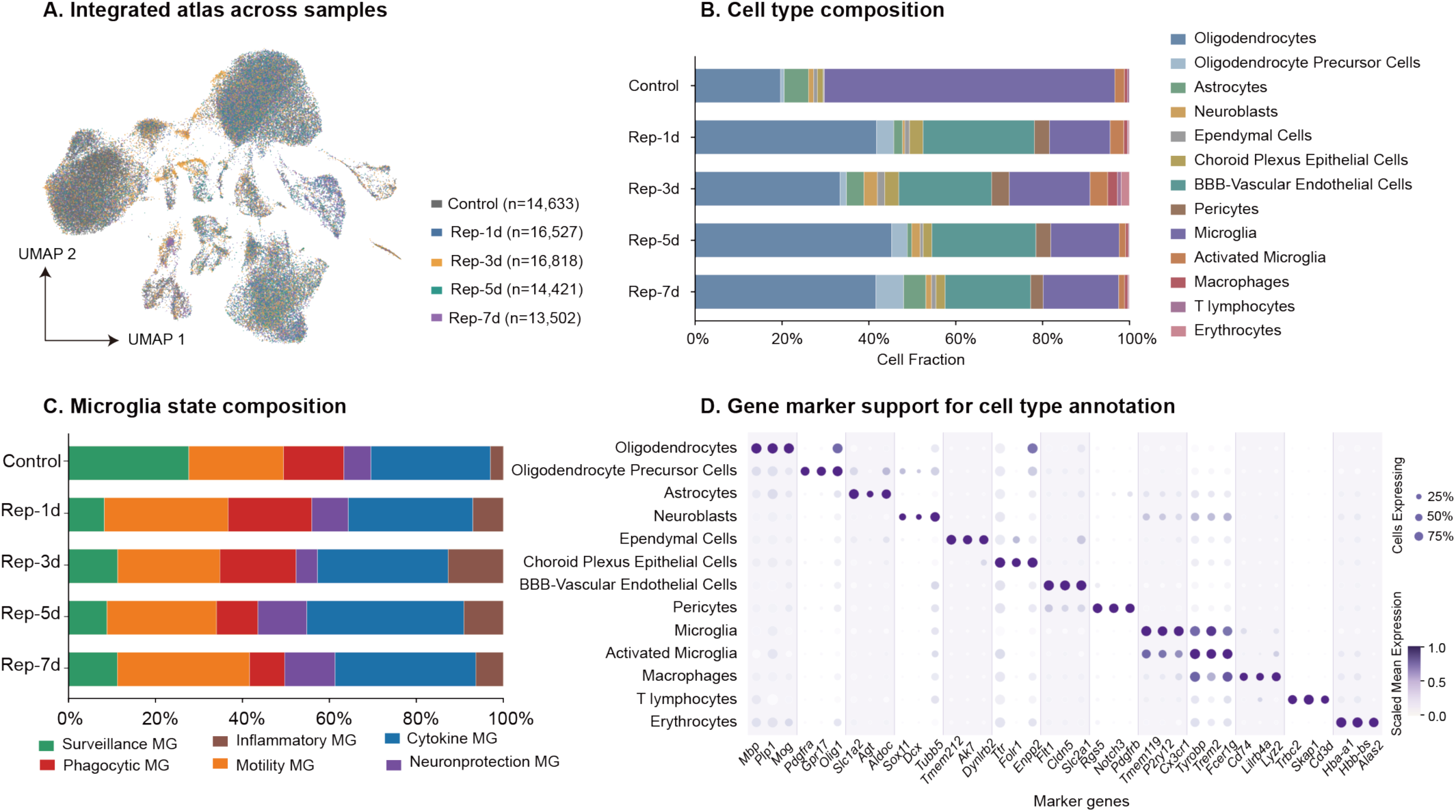
Integrated single cell atlas and cellular composition across experimental groups. **A** UMAP visualization of the integrated single cell atlas colored by sample origin. **B** Relative proportions of the 13 annotated cell types across experimental groups. **C** Relative proportions of six microglial states across groups. **D** Dot plot showing marker gene support for cell type annotation. Dot size indicates the percentage of cells expressing each marker, and color indicates scaled mean expression.

**Supplementary Fig. 11.**
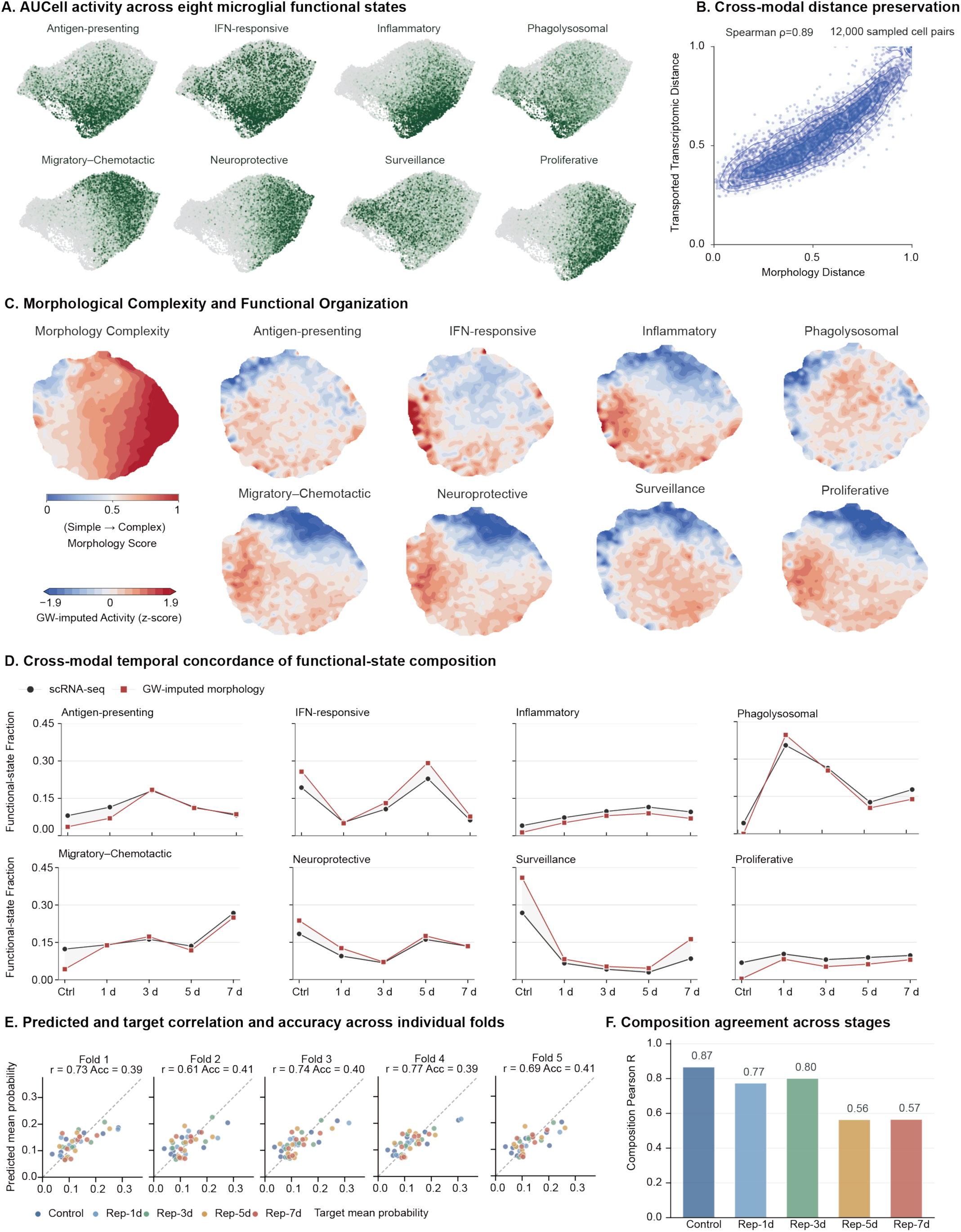
Cross modal validation of microglial morphology–function alignment and prediction. **A** Single cell UMAPs showing AUCell activity for eight microglial functional states. Darker green indicates higher AUCell activity**. B** Pairwise distance correspondence between the morphology and GW transported transcriptomic spaces at Rep-3d for 12,000 sampled cell pairs. Distances are normalized to 0–1, with points and density contours showing their joint distribution. **C** Morphological complexity and GW imputed functional activity mapped onto the morphology UMAP. The morphology score ranges from 0 (simple) to 1 (complex), and functional state maps show GW imputed activity as standardized scores. **D** Temporal trajectories of eight functional state fractions across Control and 1, 3, 5 and 7 days after reperfusion. Black circles indicate scRNA seq derived fractions, and red squares indicate morphology imputed fractions using GW-OT. **E** Correlation between target and predicted mean functional state probabilities in each cross validation fold. Points represent condition state combinations and are colored by experimental group. **F** Correlation between predicted and target functional state compositions at each reperfusion stage.

**Supplementary Fig. 12.**
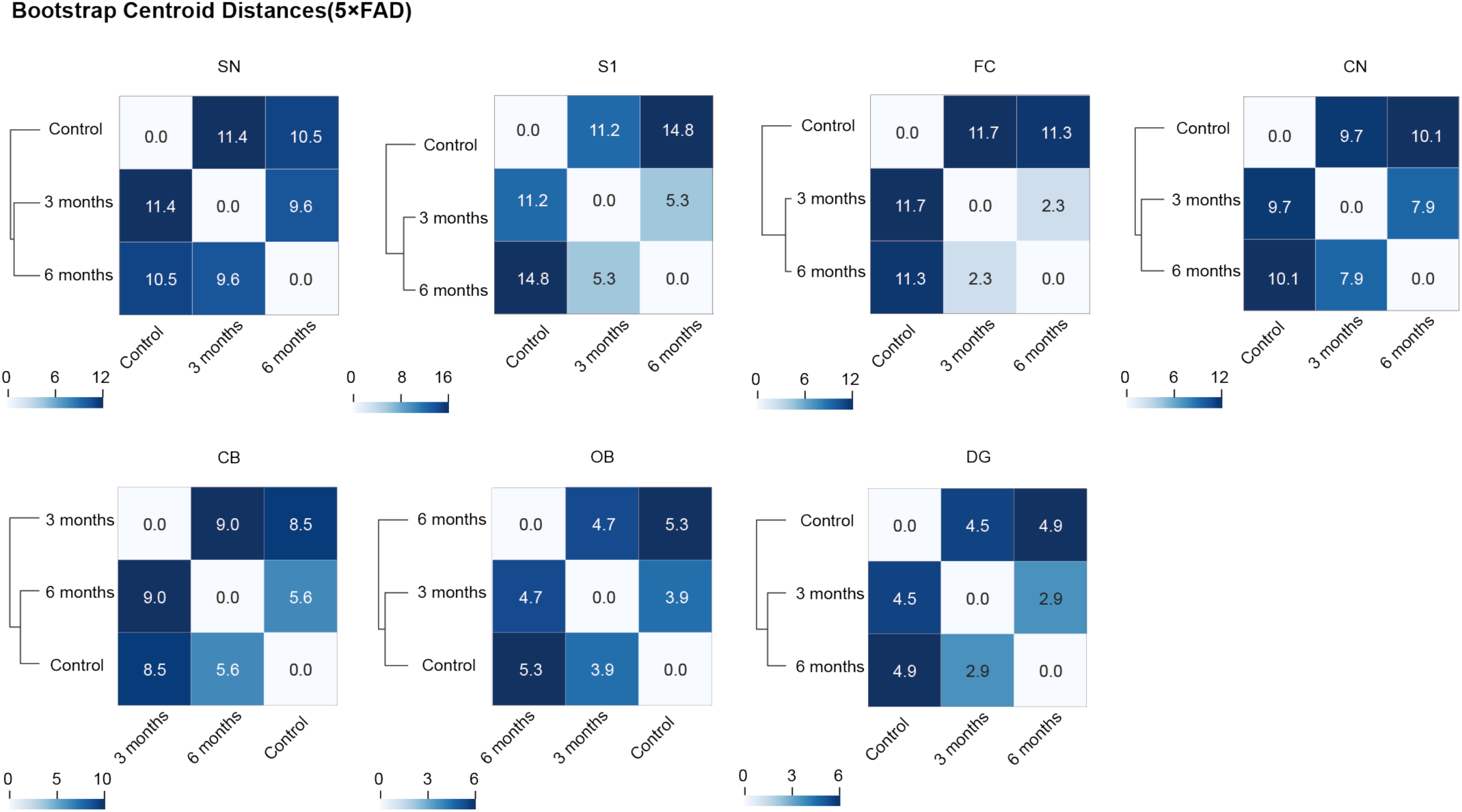
Centroid distance analysis across 5×FAD stages. Heatmaps show pairwise Euclidean distances between bootstrap centroids across Control, 3 and 6 month groups in different brain regions. Dendrograms show hierarchical relationships among groups. Cell values indicate centroid distances, with darker blue indicating greater morphological separation.

**Supplementary Fig. 13.**
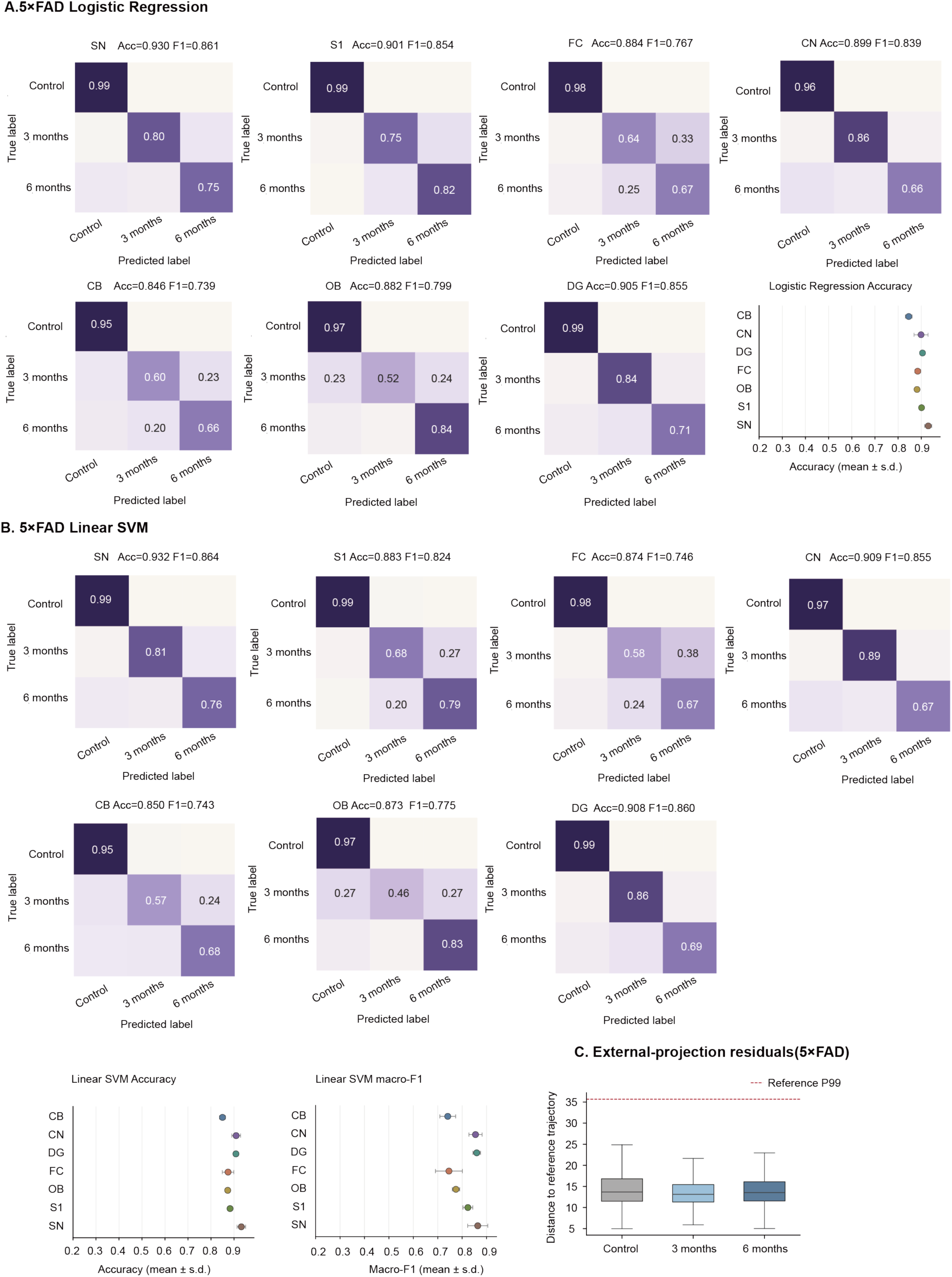
Regional separability and fixed reference projection of microglial morphology across 5×FAD stages. **A** Confusion matrices for logistic regression classification of Control, 5×FAD 3 month and 5×FAD 6 month cells within each brain region (SN, S1, FC, CN, CB, OB and DG). **B** Corresponding confusion matrices for multinomial linear support vector machine classification. **C** Distances of externally projected cells from the fixed reference trajectory across 5×FAD stages. Boxes show medians and interquartile ranges, and the red dashed line indicates the reference P99 residual threshold.

**Supplementary Fig. 14.**
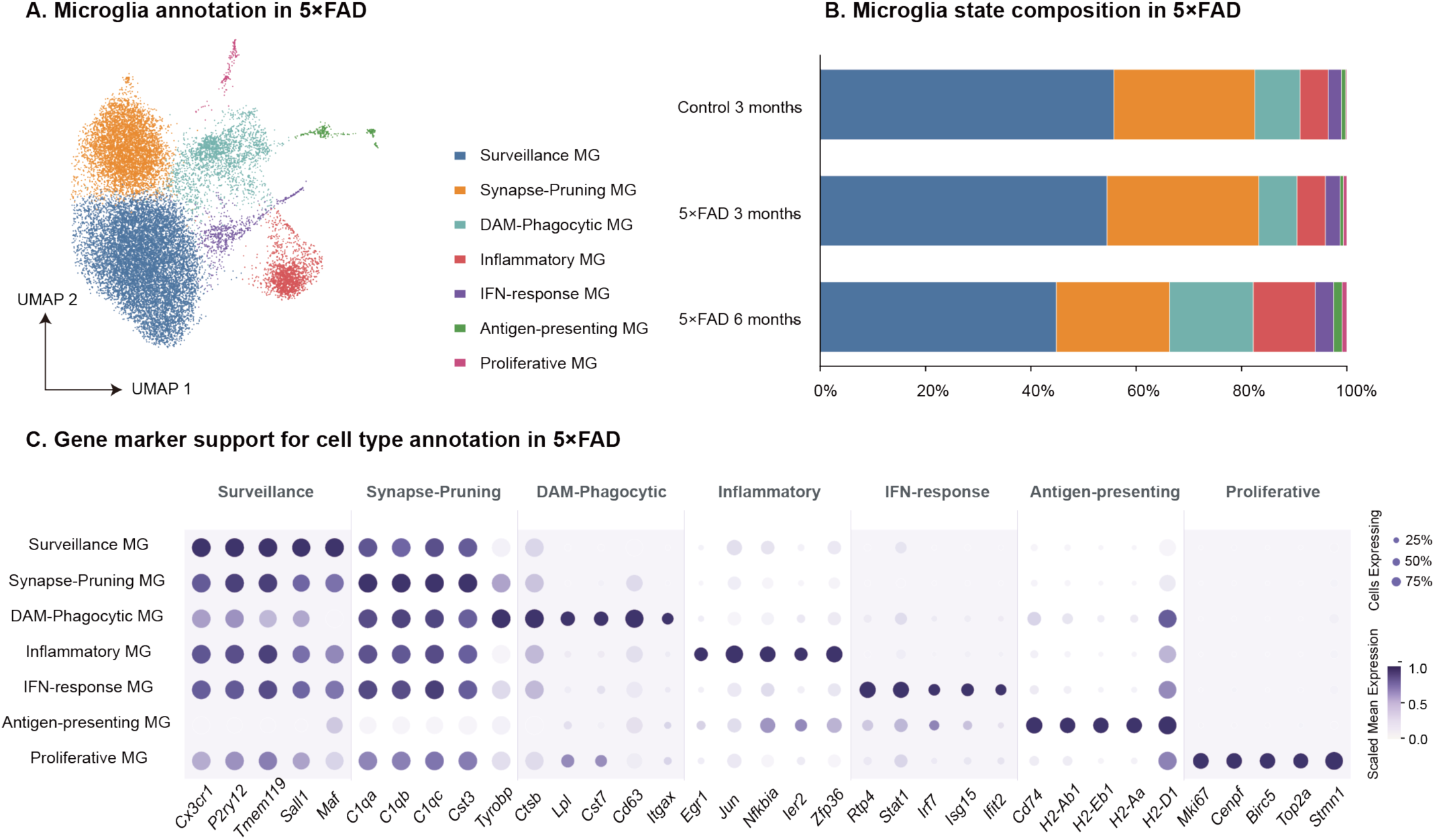
Microglial states across 5×FAD progression. **A** UMAP visualization of seven annotated microglial states. **B** Relative proportions of microglial states across Control, 5×FAD 3 month and 5×FAD 6 month groups. **C** Dot plot of marker genes supporting microglial state annotation. Dot size indicates the percentage of expressing cells, and color indicates scaled mean expression.

**Supplementary Fig. 15.**
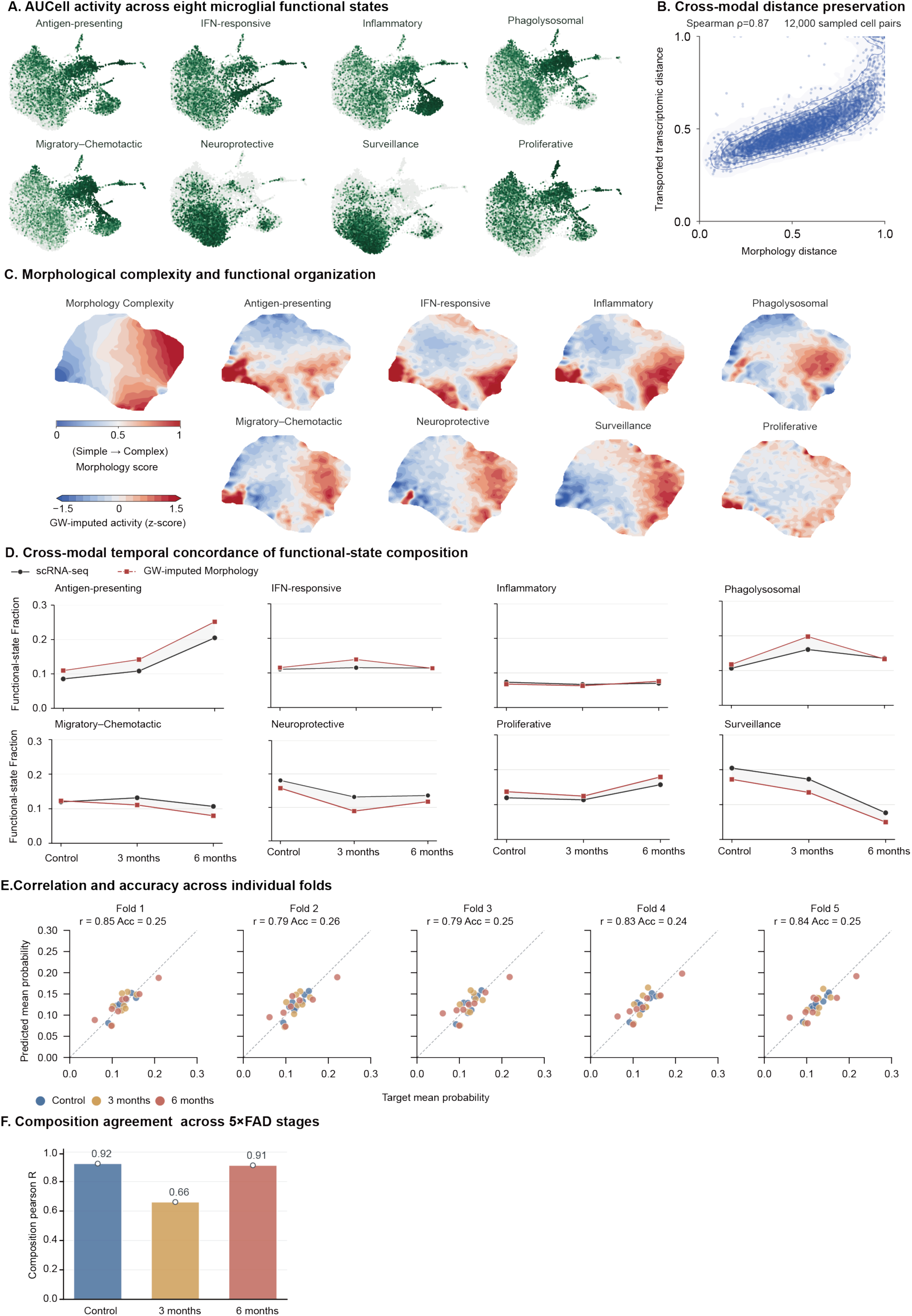
Cross modal validation of microglial morphology function alignment and prediction in 5×FAD. **A** Single cell UMAPs showing AUCell activity for eight microglial functional states. Darker green indicates higher AUCell activity. **B** Pairwise distance correspondence between the morphology and GW transported transcriptomic spaces in 5×FAD for 12,000 sampled cell pairs. Distances are normalized to 0–1, with points and density contours showing their joint distribution. **C** Morphological complexity and GW imputed functional activity mapped onto the morphology UMAP. The morphology score ranges from 0 (simple) to 1 (complex), and functional state maps show standardized GW imputed activity. **D** Temporal changes in eight functional state fractions across Control, 5×FAD 3 month and 6 month groups. Black circles indicate scRNA seq derived fractions, and red squares indicate morphology imputed fractions using GW-OT. **E** Correlation between target and predicted mean functional state probabilities in each cross validation fold. Points represent condition state combinations and are coloured by experimental group. **F** Correlation between predicted and target functional state compositions across 5×FAD stages

